# A comprehensive analysis of single-cell transcriptome network underlying gastric premalignant lesions and early gastric cancer

**DOI:** 10.1101/455121

**Authors:** Peng Zhang, Mingran Yang, Yiding Zhang, Shuai Xiao, Xinxing Lai, Aidi Tan, Shao Li

## Abstract

Intestinal-type gastric cancer is preceded by premalignant lesions including chronic atrophic gastritis (CAG) and intestinal metaplasia (IM), which are characterized as changes in cell types. In this study, for the first time, we systematically constructed a single-cell atlas for a total of 31,164 high-quality cells from gastric mucosa biopsies of patients spanning a cascade of gastric premalignant lesions and early gastric cancer (EGC) using single-cell RNA sequencing (scRNA-seq). Based on the atlas, we construct a network underlying the changes of cellular and molecular characteristics of gastric epithelial cells across different lesions. We found the conversion of gland mucous cells (GMCs) toward a more intestinal-like stem cell phenotype during metaplasia, and identified OR51E1 as a novel marker for early-malignant enteroendocrine cells. We also found that HES6 might mark a goblet cell subset that precede morphologically identifiable goblet cells in IM mucosa, potentially aiding the identification of metaplasia at the early stage. Finally, we identified a panel of EGC-related specific signature, with clinical implications for the precise diagnosis of EGC. Our study offers unparalleled insights into the human gastric cellulome in premalignant and early-malignant lesions and provides an important data resource that will facilitate studies in gastritis-induced tumourigenesis and gastric cell biology.

**Significance Statement:** Understanding cellular characteristics in gastric premalignant and malignant lesions would help us better understand the gastric cancer (GC) pathogenesis. In this paper, for the first time, we systematically constructed a single-cell transcriptome network of human premalignant gastric mucosa and early GC (EGC) and derived novel findings from it. We identified OR51E1 as a novel marker for early-malignant enteroendocrine cells and a panel of genes as the EGC-specific signature, with clinical implications for the precise diagnosis of EGC. We also found HES6 might mark a goblet cell subset that precede morphologically identifiable goblet cells in IM mucosa, potentially aiding the identification of metaplasia at the early stage. Our study provided an unprecedented data resource that will facilitate studies underlying gastritis-induced tumorigenesis.

## Introduction

According to the Correa model, chronic atrophic gastritis (CAG) and intestinal metaplasia (IM) are the main premalignant lesions of intestinal-type gastric cancer (GC)^1^. Patients with CAG or IM are at an increased risk of GC, with the estimated annual risk of GC being 0.1% per year in CAG patients and 0.25% per year for IM patients^2^. The change in cell types plays a crucial role in the cascade from premalignant lesions to the malignant lesion. For example, CAG is characterized as the loss of parietal cells while IM is defined as the emergence of intestinal-specific cell types, including goblet cells and enterocytes^3–5^. However, the full spectrum of distinct cell types and their molecular characteristics remain to be well defined in the gastric mucosa of premalignant and malignant lesions, which hampers our ability to investigate their roles in GC pathogenesis.

The cellular characteristics are crucial for identifying gastric premalignant and malignant lesions. In the normal gastric mucosa, the epithelium is constructed from a complex compendium of cell lineages including mucous cells, secretory cells and endocrine cells, which work in coordination to maintain tissue homeostasis^6,7^. The damage caused by the Helicobacter pylori infection and other risk factors to the gastric mucosa is characterized by the partial and, finally, total loss of glandular cells, thus developing as CAG^8^. During the progression of gastritis, IM develops in the mucosa with the emergence of some metaplastic cell lineages, including intestinal-specific goblet cells and enterocytes^5^. Alternatively, another type of metaplasia, termed spasmolytic polypeptide–expressing metaplasia (SPEM), is characterized as the presence of TFF2+ cells and has also been reported to show a strong association with GC.^9^. Additionally, some types of cells have been observed within the gastric mucosa of different lesions. For examples, goblet cells, the intestinal-specific cell lineage present in the IM mucosa, have also been observed in the dysplastic glands^10^. Therefore, it is important to systematically characterize cell lineages within the gastric mucosa of each lesion. However, previous studies that explored the molecular features of GC and premalignant stages using bulk sample-based experiments or mathematical models have obscured the signatures of distinct cell populations^11–17^. Alternatively, other studies that relied on predetermined markers to purify cell populations have also been limited to several specific cell types, failing to fully distinguish between cell types and to detect rare cellular populations or cells with specific states^6, 18–20^. Therefore, a comprehensive and systematic profile of the expression of individual cells in gastric premalignant and malignant mucosae is urgently needed.

Advances in single-cell mRNA sequencing (scRNA-seq) have revolutionized our ability to characterize the transcriptional state of thousands of individual cells in-depth, enabling an unbiased analysis of the spectrum of cell populations within tissues. This technology has been successfully applied to identify cell types and understand the complex subpopulations in tissues of organs such as the pancreas^21^ or lungs^22^, as well as in various cancers including melanomas and colon cancer^23,24^.

Here, we performed a scRNA-seq survey of 54,575 cells from twelve gastric mucosa biopsies from eight patients with Non-atrophic gastritis (NAG), CAG, IM and early gastric cancer (EGC), and constructed a single-cell transcriptome atlas for gastric premalignant and early-malignant lesions for the first time. Using the atlas, we construct a network for dissecting the changes of cellular and molecular characteristics of gastric epithelial cells across different lesions. We then characterized the expression profiles of two conserved cell types, gastric mucous-secreting cells and enteroendocrine cells, across different lesions. We also identified two subsets that reflecting distinct cell states within goblet cells. Finally, we systematically analyzed the expression profile of EGC cells and identified a panel of high-confidence markers for precise identification of EGC. This large-scale single cell atlas provides unprecedented insights into the cellular heterogeneity within gastric mucosa in different lesions.

## Results

### A single-cell atlas of premalignant gastric mucosae and early gastric cancer

To characterize the single-cell profile of gastric mucosa in premalignant and early-malignant lesions, a total of twelve biopsies, including two NAG biopsies, three CAG biopsies, six IM biopsies and one EGC biopsy, were taken from eight patients, which span the cascade from gastritis to early gastric cancer (Figure 1a). Individual 2mm biopsies were separated into two equal parts to provide tissues for scRNA-seq and histological detection, respectively. The pathological grade of each biopsy was determined immunohistologically, as shown in Supplementary figure 1, and was assessed independently by two trained pathologists using the updated Sydney classification for IM grading and the revised Vienna classification for gastric dysplasia ^25,26^ (Methods, Supplementary table 1). Each biopsy with IM was further classified as mild or severe, based on hematoxylin and eosin (H&E) staining and MUC2-based immunofluorescence (IF) staining (Supplementary figure 2). The clinical characteristics of these participants, including *H.pylori* infection and clinical symptoms, were recorded at recruitment in the Supplementary table 2. We also collected the endoscopic image of each biopsy site and tongue images for each patients (Supplementary figure 3). For each biopsy, we isolated single cells without prior selection for cell types and utilized the 10x Chromium platform to generate RNA-seq data. After removing low-quality cells (Methods), a total of 31,164 cells that passed the quality control were retained for subsequent analysis, which yielded a median of 1268 detected genes per cell. The number of cells from each biopsy was provided in the Supplementary table 3.

**Figure 1.**
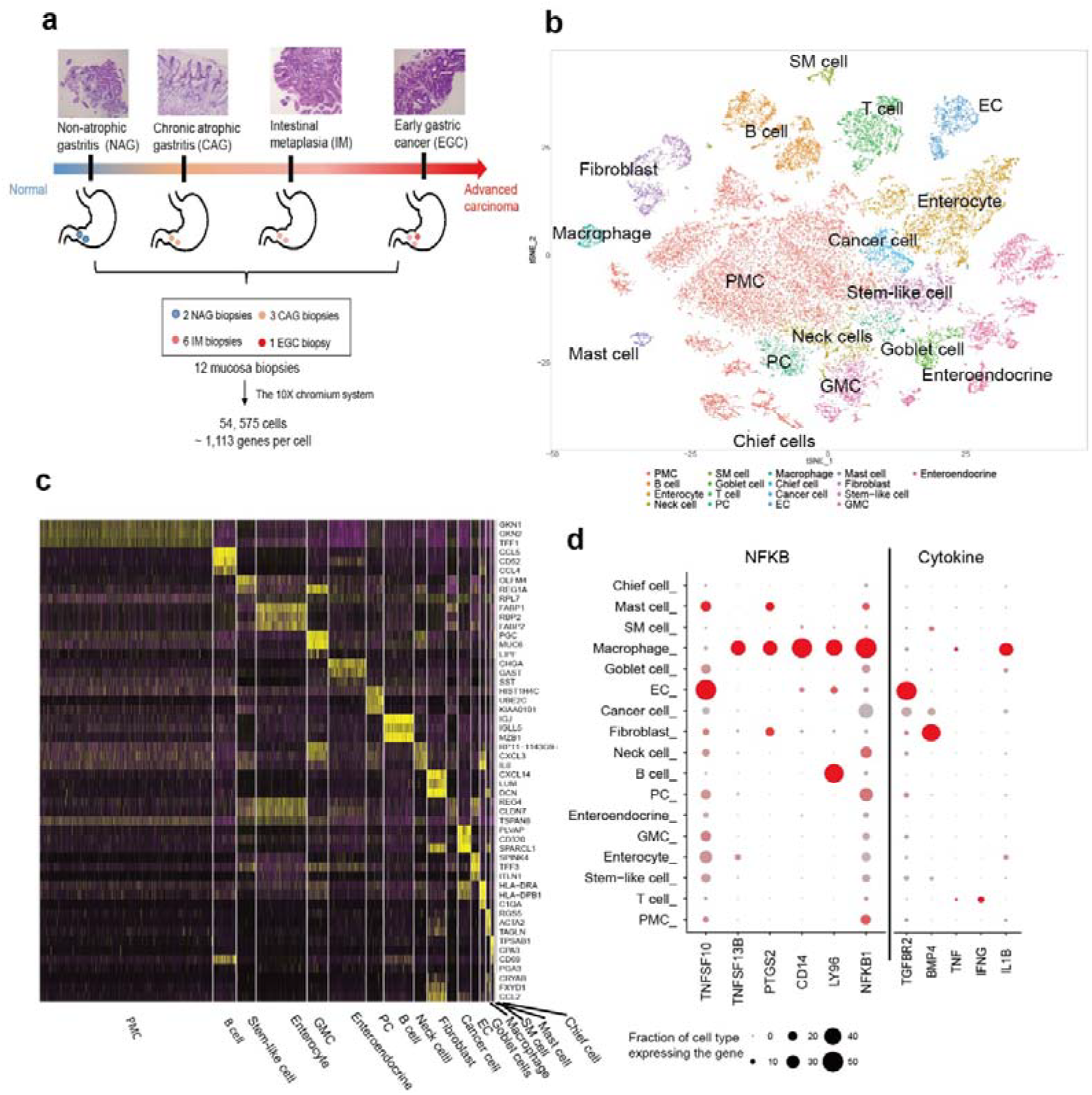
A single-cell atlas of gastric premalignant and early-malignant mucosae. **a.** Schematic diagram highlighting the experimental workflow for the whole study. Twelve mucosa biopsies were collected from eight patients with non-atrophic gastritis (NAG, 2 biopsies), chronic atrophic gastritis (CAG, 3 biopsies), intestinal metaplasia (IM, 6 biopsies) and early gastric cancer (EGC, 1 biopsy). The transcriptome of single cells was sequenced using the 10X chromium system. **b.** The t-SNE plot of 31,164 high-quality cells to visualize cell type clusters based on the expression of known marker genes. GMC, gland mucous cell; PC, proliferative cell; PMC, pit mucous cell; SM cell, smooth muscle cell. **c.** Cell-type markers. The relative expression level of genes across cells is shown, sort by cell type. Cell-type marker genes were identified in an unbiased fashion (Wilcoxon rank sum test, FDR < 0.01 and fold change > 1.5) and only top three were shown in the figure. **d.** Dot plot of representative genes in the NFKB signaling pathway and cytokines mapped onto cell types.

To identify distinct cell populations based on gene expression patterns, we performed dimensionality reduction and unsupervised cell clustering using methods implemented in the Seurat software suite ^27^, followed by removing batch effects among multiple samples (Methods). As shown using t-distribution stochastic neighbor embedding (t-SNE)^28^, profiles along the cascade from NAG, CAG, IM, to EAG were derived and a total of 17 main cell clusters were finally identified (Figure 1b, Supplementary figure 4 and 5, Methods), which we defined as the single-cell transcriptome atlas of human gastric mucosa in premalignant and early-malignant lesions. Based on the expression of known marker (Supplementary table 4), we found the atlas mainly comprised epithelial cells (EPCAM) and non-epithelial cells (VIM) (Supplementary figure 6). The epithelial cells mainly consisted of gastric cells, including gland mucous cells (MUC6), pit mucous cells (MUC5AC), chief cells (PGA4 and LIPF) and enteroendocrine cells (CHGA and CHGB). We annotated one epithelial cell cluster as proliferative cells due to canonical cell cycle markers, such as MKI67 and BRIC5. We also annotated one epithelial cluster that almost uniquely expressed in the EGC biopsy and significantly up-regulated tumor markers (CEACAM5 and CEACAM6) as cancer cells. One of the epithelial cell clusters was annotated as stem cell due to the elevated expression of the stem cell marker OLFM4 ^29^. As expected, the epithelial cells also comprised goblet cells (MUC2) and enterocytes (FABP1 and APOA1), which was agreeable with the cellular characteristics of IM. The non-epithelial cell lineages comprised T cells (CD2 and CD3D), B cells (CD79A), macrophages (CSF1R and CD68), fibroblasts (DCN and PDPN), smooth muscle cells (ACTA2), endothelial cells (VWF and ENG), and mast cells (TPSAB1). Apart from canonical cell type markers, we also identified additional genes that strongly and specifically marked each major cell population (Figure 1c, Supplementary table 5). Kyoto Encyclopedia of Genes and Genomes (KEGG) pathway and Gene Ontology (GO) enrichment analysis of genes with restricted expression in each major cell cluster supported predicted identities of cell populations. For example, enterocytes mainly involved in metabolic pathways (FDR < 2E-50) and fibroblast markers mainly involved in terms about extracellular matrix organization (FDR < 1.5E-12). We also investigated the cell heterogeneity by analyzing of the cell-to-cell correlations. We observed a slight increase in the correlation between epithelial cells from the same sample, compared to that between non-epithelial cells. The latter showed an obvious increase in the correlation between cells in the same lesion (Supplementary figure 7). Thus, we identified a broad range of cell types from gastric mucosa samples in the premalignant and early-malignant lesions.

Next, we plotted the cell-of-origin for some mediators in the cytokines and NFKB signaling pathway, respectively, which were known for involving gastritis-induced gastric tumorigenesis^30^ (Figure 1d). For example, we found that macrophages might be a dominant source of IL1B, whose genotype was reported to confer greater risk of gastric cancer^31^, and PTGS2, which play a crucial role in mediating the inflammatory process through activation of NF-κB^32^, suggesting macrophages in the inflammatory microenvironment act as a pivotal role to promote gastric tumorigenesis. In addition, BMP4, which is broadly reported to regulate gastric epithelial cell development and proliferation^33^, was found to express uniquely restricted to fibroblasts, suggesting a link between activation of fibroblasts and gastric epithelial cell proliferation. Thus, we could conclude that our single-cell atlas provided unbiased insights for the cellular source of known mediators underlying gastritis-induced gastric cancer.

### Construction of single cell-based network for characterizing gastric epithelial cells across different lesions

We first focused on changes of epithelial cells in different lesions along the cascade from gastritis to gastric cancer. Through clustering analysis, we identified certain epithelial cell types, such as PMC, GMC, and endocrine cells (Figure 2a). Among them, we found some ‘conservative’ cell types that presented across different lesions, such as gastric mucous cells, proliferative cells, stem-like cells and endocrine cells, whereas others emerged in certain lesions, such as goblet cells and enterocytes in the IM lesion and cancer cells in the EGC lesion (Figure 2b). In particular, we observed that the proportion of gastric highly differentiated cell types, including pit mucous cells (PMCs), decreased along the cascade. In contrast, the proportion of stem-like cells increased significantly in the IM lesion, and reached the highest in the EGC lesion (Supplementary figure 8).

Further, we systematically profiled the spectrum of gene expression of epithelial cells across different lesions. Using the Wilcoxon rank sum test, we identified differential expression genes (DEGs) for each cell type in each lesion (FDR < 0.01, fold.change > 1.5, Supplementary table 6). Then, we merged DEGs of multiple cell types in the same lesion as the lesion-related signature and observed a clear distinction of cells in mucosa at distinct lesions with those signatures (Figure 2c). Interestingly, we observed that these lesion-related signatures showed significant overlap with those derived from the bulk transcriptome dataset (GSE2669), as well as high-risk genes inferred for gastritis or gastric cancer by the state-of-art disease-gene prediction algorithm^34^ (Figure 2d). We then identified pathways that significantly enriched by signatures for each lesion (FDR < 0.05, Supplementary table 6) and found that most of them were previously well-documented involving in the gastritis and gastric cancer^35^, such as TNF signaling pathway and mineral absorption in the CAG lesion, the metabolism-related pathways in the IM lesion, and cell proliferation-related pathways in the EGC lesion.

**Figure 2.**
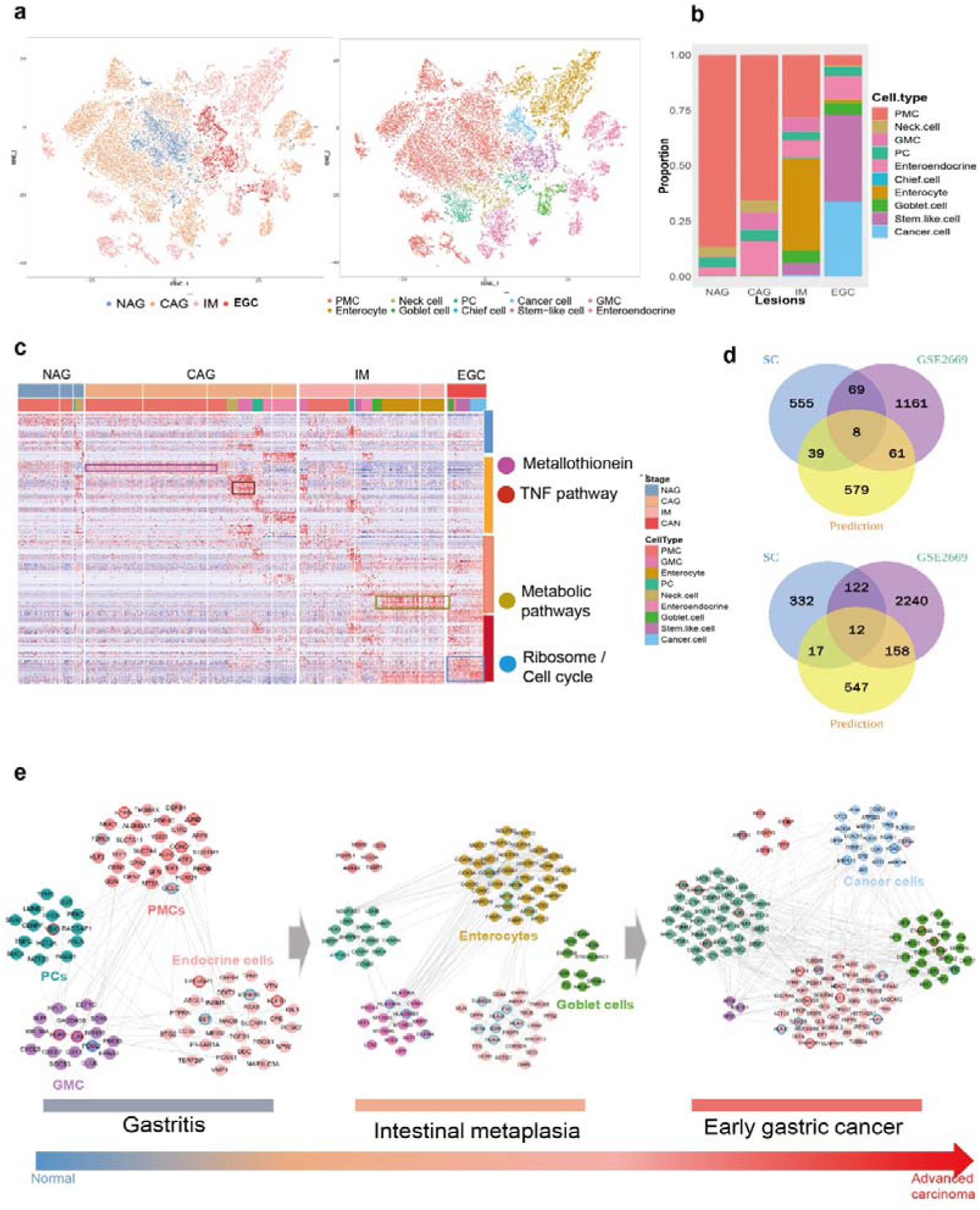
The single-cell transcriptomes of epithelial cells in the cascade from gastritis to EGC. **a.** t-SNE plot of 24,223 re-clustered epithelial cells from the gastric mucosa by lesions (non-atrophic gastritis (NAG, 3,306 cells), chronic atrophic gastritis (CAG, 10,966 cells), intestinal metaplasia (IM, 7,815 cells) and early gastric cancer (EGC, 2,136 cells) and cell type (right). **b.** The proportion of epithelial cell types across multiple lesions. The cell types were annotated based on figure 2a. **c.** The heatmap showing expression patterns of multiple epithelial cell types in premalignant gastric mucosa and early gastric cancer. Genes involved in the heatmap were significantly up-regulated genes (FDR < 0.01, fold change > 1.5, Wilcoxon rank sum test) for each cell type in each lesion. Pathways that mostly enriched for the gene signature in each lesion were denoted at the right side. **d.** Venn diagram for the gene signatures identified by different evidences including the single-cell atlas, bulk transcriptome dataset (GSE2669) and network-based prediction^34^, in multiple lesions (upper panel, gastritis; lower panel, gastric cancer). SC, single cell. **e.** The single-cell transcriptome network underlying gastric premalignant and early malignant lesions. Nodes with different color represented different cell types and those with cycle represented high-risk genes inferred by CIPHER for gastritis or gastric cancer (red, gastritis; green, gastric cancer).

It should be noted that we could further dissect the cell types in which lesion-related signatures were preferentially expressed (Figure 2c). Although both were identified as the molecular characteristic for the CAG lesion, genes involved in mineral absorption preferentially expressed in the PMCs while ones involved in the TNF signaling pathway, including CXCL2, CXLC3, tended to express in the GMCs. Since CXCL2 and CXCL3 are potent stimulators of neutrophil activation and function in neutrophil attracting, we could speculate that neutrophils appeared to be recruited around the GMCs under the condition of chronic gastritis. Metabolic pathways, the main molecular characteristics in the IM lesion, were found to significantly upregulated inenterocytes. In the EGC lesion, cell proliferation-related genes are highly expressed in almost all cell types in EGC, suggesting that cell proliferation is a common feature of gastric mucosal epithelial cells in the EGC lesion.

To systematically contextualize both molecular and cellular shifts in the cascade from gastritis to EGC, a network underlying the cascade from gastritis to gastric cancer was finally constructed by linking signature genes for each cell type in each lesion through the protein-protein interactions (Figure 2e, Methods). In the network, nodes with different colors represent different cell types in which signature genes preferentially express. Thus, our single cell-based network provided the systematic spectrum of changes of human gastric epithelial cells across premalignant and EGC lesions, with both molecular and cellular aspects.

### Characterization of the single cell expression profiles for gastric mucous-secreting cell lineages across different lesions

From the single-cell-based network in the figure 2e, the gastric mucous-secreting cell is the ‘conserved’ cell type across lesions (Figure 3a), and is mainly composed of MUC5AC-expressing PMCs and MUC6-expressing GMCs (Figure 3b and c). We found these two cell types showed distinct expression patterns where PMCs mainly expressed genes involved in actin cytoskeleton while GMCs mainly expressed immune-related genes (Figure 3d, Supplementary table 7). For the PMCs, We found that the molecular hallmarks previously implicated for each lesion were also reflected within the PMCs in different lesions (Supplementary figure 9). For example, the CAG-related markers, GAST and PGC, showed significant up-regulation in the PMCs in the CAG lesion and the GC-related genes, REG4 and CDH17, showed significant up-regulation in the PMCs in EGC lesion (Figure 3e). Interestingly, we also found that the enterocyte-specific gene, FABP1, was also up-regulated in the PMCs in the IM lesion (Figure 3e). Therefore, it revealed that a dysregulation of fatty-acid metabolism would not only occur in the intestinal cells but also in the gastric cells during metaplasia.

**Figure 3.**
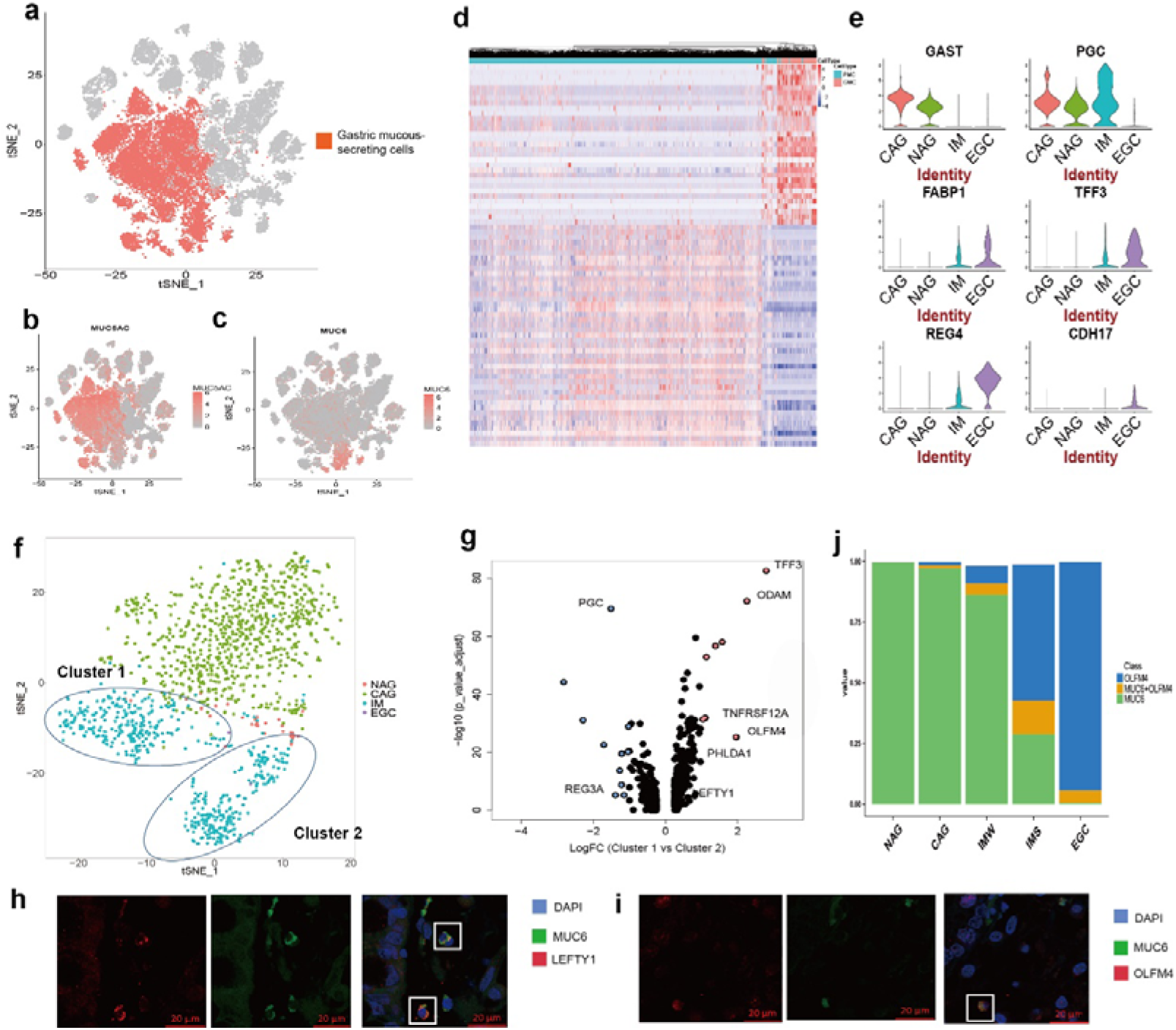
The scRNA profiles for gastric mucous-secreting cell lineages across different lesions. **a.** The t-SNE plot that showed the distribution of gastric mucous-secreting cell lineages (red, n = 14,481) in the atlas, with marked by the expression of marker genes, MUC5AC (**b**) and MUC6 (**c**), respectively (red = high, gray = low). **d.** The heatmap showing expression of selected functionally relevant genes that are differentially expressed between the two sub-clusterstypes of gastric mucous-secreting cell lineages, GMC and PMC (Fold change > 1.5, FDR < 0.01). **e.** Violin plots display the distribution of expression of known lesion-specific genes (CAG, upper; IM, middle; EGC, bottom) in PMCs in diverse lesions. **f.** The t-SNE plot showing GMCs across diverse lesions. **g.** Volcano plot showing fold change and P value of genes differentially expressed between two globle cell sub-clusters (FDR < 0.01). Genes with the absolute value of the fold change of expression between two sub-clusters larger than 1.5 were colored red (upregulated in cluster 1) or blue (upregulated in cluster 2). Selected highly significant genes are labelled. Genes labeled red are ones show elevated expression in Cluster 1 while genes labeled blue are ones show the elevated expression in Cluster 2. **h, i.** Double immunofluorescence staining images of the pair of MUC6 and LEFTY1 (**h**), and the pair of MUC6 and OLFM (**i**), in the IM biopsy. Scale bar, 20 μM; nuclei (DAPI) in blue. **j.** Bar chart showing the relative percent of cells expressing MUC6 or OLFM4 alone or in combinations (MUC6+OLFM4) across different lesions.

Interestingly, we observed the high cellular heterogeneity in the gastric gland cells, marked by MUC6, in the IM lesion. Projected into the 2-dimensional PCA-space, these MUC6-expressing gland cells were clearly divided into two subclusters (denoted as Cluster 1 and Cluster 2, respectively, Figure 3f). The expression signature of the Cluster 1 was enriched for immune- and antimicrobial-related genes, which was agreeable with molecular characteristics in the normal antral gland cells while the expression signature of the Cluster 2 mainly consisted of intestinal stem cell or development-related genes, including OLFM4, PHLDA1, ODAM, and LEFTY1 (Figure 3g). The co-expression pattern of MUC6 and OLFM4, as well as LEFTY1, was confirmed by the immunofluorescence (IF) staining on the same IM resection specimen (Figure 3h and i). Of note, OLFM4 has been reported to marks intestinal stem cells in both normal and metaplastic contexts. Therefore, we speculated that GMCs tended to acquire the intestinal stem cell phenotype in the IM lesion.

Next, we extended to examine the change of the co-expression pattern of OLFM4 and MUC6 in individual GMCs across diverse lesions. Using the threshold at the tenth centile to include 90% of cells with at least one transcript detected from each gene of interest, we observed gradual increase of the proportion of OLFM4-expressing GMC cells with further intestinalization of gastric mucosa. As shown in Figure 3j, GMCs with expressing the OLFM4 were rarely detected in the CAG lesion (0.4%), while the number of them (remarkably) increase in the wild IM lesion (8%) and reach a peak in the severe IM lesion (26%). In the EGC lesion, the GMC disappeared and the proportion of OLFM4-expressing cells reach the peak, which have been confirmed by previous immunohistochemistry studies^36^. We further assessed the co-expression of MUC6 and OLFM4 within the GMCs in the IM lesion, and found that the expression of OLFM4 displayed a significant negative correlation with that of MUC6 (p = 0.003, Supplementary figure 10). Additionally, the OLFM4 was mainly expressed in the GMCs (24.2 %), compared to surface pit cells (3.5%). These data suggest the conversion of GMCs toward a more intestinal-like stem cell phenotype is the crucial cellular characteristics for gastric intestinal metaplasia, as well as gastric tumorigenesis.

### Characterization of the single-cell expression profiles for enteroendocrine cell lineages across different lesions

From the single-cell-based network shown in the figure 2e, the enteroendocrine cell is another ‘conserved’ cell type across different lesions. Enteroendocrine cells secrete hormones and peptides that play important roles in digestion, inflammatory processes, and energy metabolism^37–40^ and they might be involved in the process underlying inflammation-induced tumourigenesis^41,42^. However, the full spectrum of enteroendocrine cell composition and heterogeneity in each premalignant lesion and early-malignant lesion remained to be determined.

To characterize the expression profiles of enteroendocrine cells within the gastric mucosa in diverse lesions, we focused on the ‘enteroendocrine cell’ cluster, which includes a total of 1760 cells (Figure 4a). We firstly observed the high heterogeneity within the cell cluster and a total of eight sub-clusters were derived by re-clustering these cells, as shown in Figure 4b. We then examined the expression distribution of canonical enteroendocrine cell markers in different samples. We observed that gastric endocrine cell markers were mainly expressed in the gastritis lesion, and the expression level decreased in the progression to IM (Figure 4c). For example, GAST, the G cell marker, showed significantly down-regulatedin the biopsies with IM, compared those with CAG (FDR < 1e-5, willcox test). In contrast, intestinal endocrine cell markers were mainly expressed in the IM lesion and minimally expressed in the gastritis lesion. Interestingly, we observed that only 12.4% of EGC endocrine cells expressed intestinal or gastric endocrine canonical markers. It revealed that endocrine cells in different lesions might be dominated by different enteroendocrine cell subtypes.

**Figure 4.**
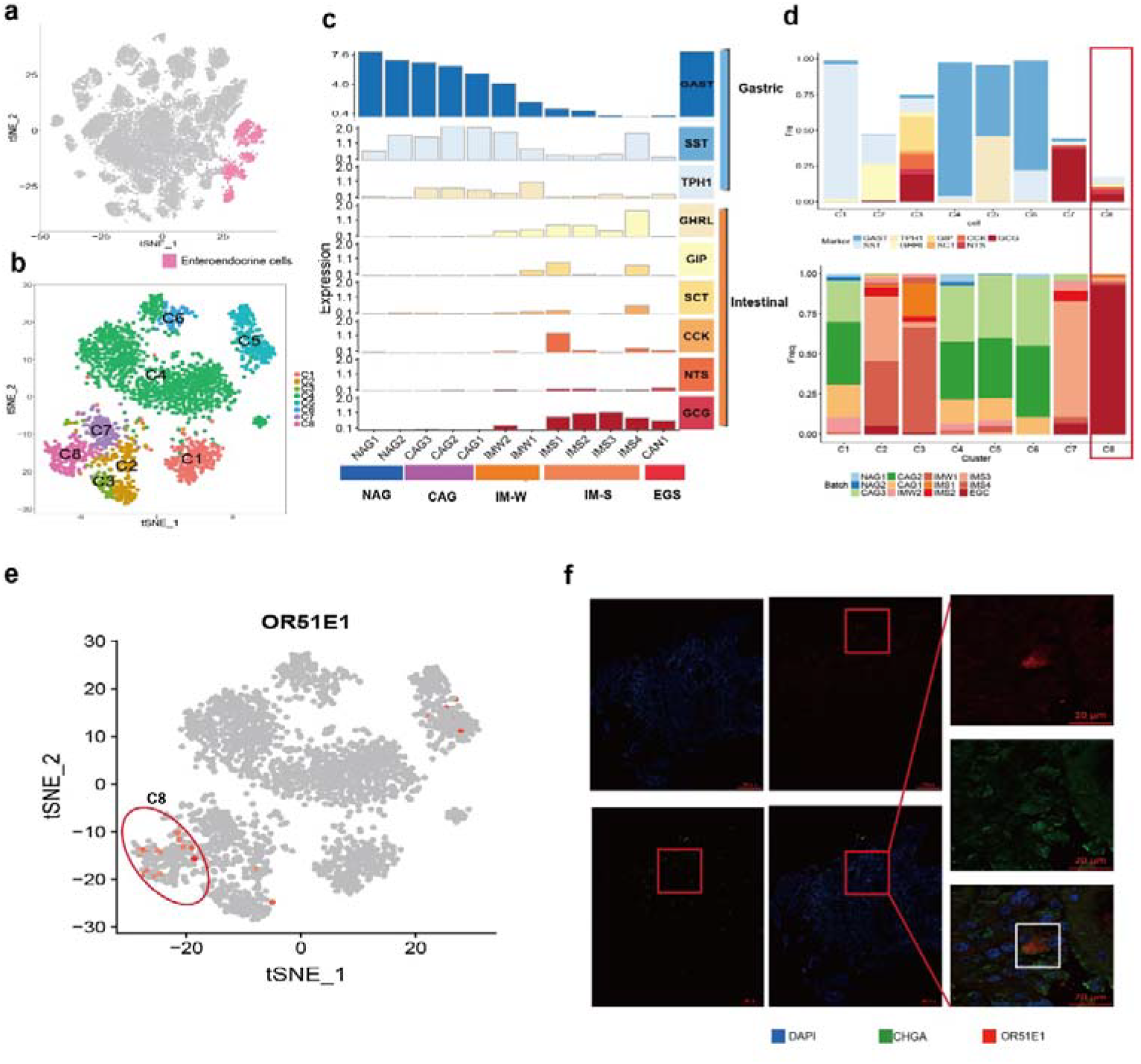
The scRNA profiles for enteroendocrine cell lineages across different lesions. **a.** The t-SNE plot that showed the distribution of enteroendocrine cell lineages (pink, n= 1,760 cells) within the atlas. **b.** Enteroendocrine cell populations were re-clustered into 8 subclusters (colour coding). **c.** The distribution of the mean expression of intestinal and gastric endocrine cell canonical markers in each sample. Gastric endocrine cell markers are displayed in blue while intestinal endocrine cell markers in red. IM-W, wild IM; IM-S, severe IM. **d.** Fraction of cells expressing canonical markers (upper) and cells at different lesions (lower) in each cluster. Y axis represents the proportion of cells expressing canonical markers while X axis represents the identified 8 clusters. **e.** The t-SNE plot shows the expression of putative EGC endocrine cell marker, OR51E1, in the whole endocrine cell lineage (red = high, gray = low). The cells within the red circle is the EGC endocrine cells. **f.** Double immunofluorescence staining images of OR51E1 and CHGA in the EGC biopsy from the Patient Px. The representative view of co-staining of OR51E1 and CHGA is shown in the enlarged images on the right. Scale bars are 200μm and 20μm in the enlarged images.

To define putative enteroendocrine cell subtypes in each lesion, we quantified the proportion of cells expressing canonical enteroendocrine cell markers within each cluster shown in figure 4b. It was found that distinct endocrine cell markers were expressed in the same cluster (Figure 4d), which is in agreement with previous results from colon epithelial samples^43^. We annotated these cell clusters with the dominate expression pattern of canonical cell markers (Methods). For example, the cluster 2 was annotated as DA/X since makers for the D cell (SST) and the A/X cell (GHRL) were expressed in up to 33% and 40%, respectively, of cells in this cluster. Collectively, we profiled the cellular composition and marker gene expression patterns for endocrine cells in both premalignant and early-malignant lesions.

Of note, we observed EGC endocrine cells showed a distinct expression pattern, where small number of cells expressed canonical enteroendocrine cell markers (Figure 4c, d). Thus, we next determined novel genes marking the EGC endocrine cell cluster. By comparing the expression profiles of EGC endocrine cells to those of the other cell lineages, we found a list of genes that were uniquely upregulated in the EGC endocrine cells. Among them, OR51E1 ranked top in the result list (FDR < 1e-10, Figure 4e), indicating that OR51E1 could be a potential marker for EGC endocrine cells. Although OR51E1 has been reported to be preferentially expressed by neuroendocrine carcinoma and enterochromaffin cells^44,45^, its role in EGC endocrine cells has not yet been demonstrated before. Additionally, we analyzed expression of OR51E1 and CHGA, the canonical endocrine cell marker, by immunofluorescence staining of IM and EGC samples, respectively. Expectedly, we observed that OR51E1 expression was detected in EGC sample (Figure 4f), but not in IM samples (data not shown). In the EGC sample, OR51E1 was generally co-expressed with CHGA (Figure 4f). Thus, OR51E1 might be identified as a new marker for the EGC endocrine cell lineage.

### Goblet cells in the IM lesion showed two distinct expression patterns

Goblet cells, emerging in the IM lesion from the network shown in figure 2e, are a requirement for the pathologic diagnosis of intestinal metaplasia of the stomach in clinical practice^46^. In our study, a total of 565 cells were classified as the ‘goblet cell’ cluster (Figure 5a) where some goblet cell-related known markers, including MUC2, SPINK4 and ITLN1^47,48^, showed most significant up-regulation (FDR < 0.01; Supplementary table 5; Supplementary figure 11). Consistent with the classification of IM, the proportion of goblet cells in severe IM (IMS) biopsies was significantly higher than that of goblet cells in the wild IM (IMW) biopsies (p < 0.01, Supplementary figure 12).

**Figure 5.**
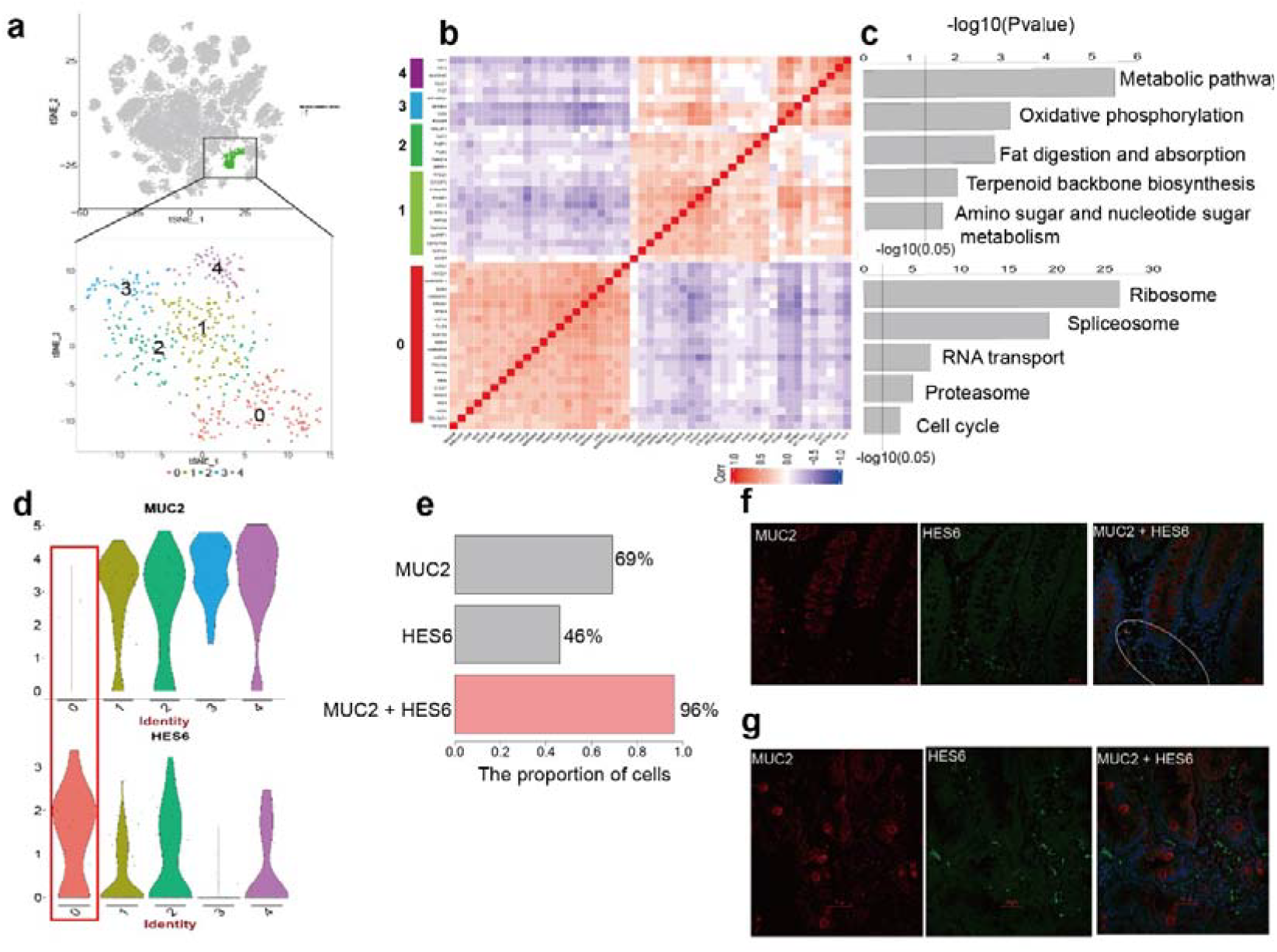
The cellular heterogeneity within goblet cell lineages in the IM lesion. **a.** The t-SNE plot that showed the distribution of the ‘goblet cell’ cluster (green, n = 565) in the atlas (upper panel) and the enlarged t-SNE plot in the lower panel showed the 5 sub-clusters goblet cell populations were re-clustered into (colour coding). **b.** The heatmap showing main signatures for the two goblet subtypes and their corresponding five most enriched pathways (FDR < 0.01) (**c**). **d.** Violin plots display the distribution of expression of MUC2 (upper panel) and HES6 (lower panel) in goblet sub-clusters. The cluster 0 is highlighted with a red rectangle. **e.** Bar chart showing the percentage of cells in the ‘goblet cell’ cluster expressing MUC2, or HES6 alone or in combination. **f.** Double immunofluorescence staining images of MUC2 (red) and HES6 (green) in the resected normal colon specimen (blue stain is DAPI). Scale bar, 50μm. **g.** Double immunofluorescence staining images of HES6 and MUC2 in the IM biopsy. Scale bar, 50μm.

Intriguingly, we also observed the high heterogeneity within goblet cells (Figure 5b). Through re-clustering these goblet cells into five sub-clusters and analyzing the co-expression of their marker genes. We observed that there existed two main patterns where one (P1), consisting of the cluster 1, 2, 3, and 4, was characterized as up-regulation of genes involved in metabolism-related pathways while another (P2), consisting of the cluster 0, was characterized as up-regulation of genes involved in cell proliferation (Figure 5c, Supplementary table 8). In details, the pattern P1 consisted of some goblet cell-related canonical markers, including MUC2, SPINK4, and ZG16, whereas the pattern P2 consisted of cell cycle-related genes, including TUBA1A. TUBB, CDK4 and HDAC. Moreover, we found that the transcription factor SOX4, with regulatory roles in stem cell maintenance and differentiation in intestinal epithelium^49^, showed the most up-regulation in the pattern P2 (FDR < 1E-15, fold change = 2.3, Supplementary table 8). Thus, it suggested that the expression pattern P2 might mark stages of goblet cell development in vivo.

Herein, we additionally identified some genes not yet implicated in goblet cells, including the Hes Family BHLH Transcription Factor 6 (HES6). HES6 is the target genes of Notch signaling, which is known to regulate the proliferation and differentiation of intestinal stem and progenitor cells. We found that HES6 transcriptionally co-expressed with SOX4 in more than half of goblet cells from IM samples (p = 7.67e-09, Supplementary figure 13) and expressed uniquely in the goblet cell cluster in the atlas (p < 1E-16, Supplementary figure 14). Moreover, we also found that HES6 exhibited mutually exclusive expression with the goblet cell-related canonical marker, MUC2, within goblet cells. HES6 was mainly expressed in the cluster 0 where MUC2 was rarely expressed (Figure 5d). Specifically, quantification of cells expressing MUC2 and/or HES6 indicated that only 69% of goblet cells expressed MUC2 alone whereas the proportion of goblet cells expressing MUC2 or HES6 reached 96% (Figure 5e).

To test this, we analyzed expression of HES6 and MUC2 by immunofluorescence staining of human colon and IM samples, respectively. In the colon sample, it was observed that HES6 was present around the crypt base, where intestinal undifferentiated or stem cells occur, while MUC2 was present in cells toward the centre or top of the crypts, where differentiated cells are found (Figure 5f). We also observed a small number cells co-expressing MUC2 and HES6. In the IM sample, we observed that most HES6-expressing cells were present around MUC2-expressing cells, many of which were located at the same metaplastic glands. Moreover, we also consistently observed cells expressing HES6 without MUC2 in isolated glands (Figure 5g). Therefore, we speculated that HES6 might play a novel role in marking an earlier stage of goblet cell differentiation and be used to mark cells with some goblet cell characteristics that are not yet morphologically identifiable as goblet cells, with potential in identifying high-risk non-IM gastritis patients in clinical practise.

### Single-cell transcriptional profiles enabled to identify signature specifically marking early gastric cancer cells

EGC is a lesion confined to mucosa and submucosa, irrespective of lymph node involvement^50^ and confers a survival rate of greater than 90% in 5 years, which renders it key to identify potential markers for early detection of gastric cancer. Herein, we focused on the putative ‘cancer cell’ cluster, emerging in the EGC lesion from the network shown in Figure 2e, and characterized its expression profile (Figure 6a), with the aim to identify EGC-related gene signatures.

**Figure 6.**
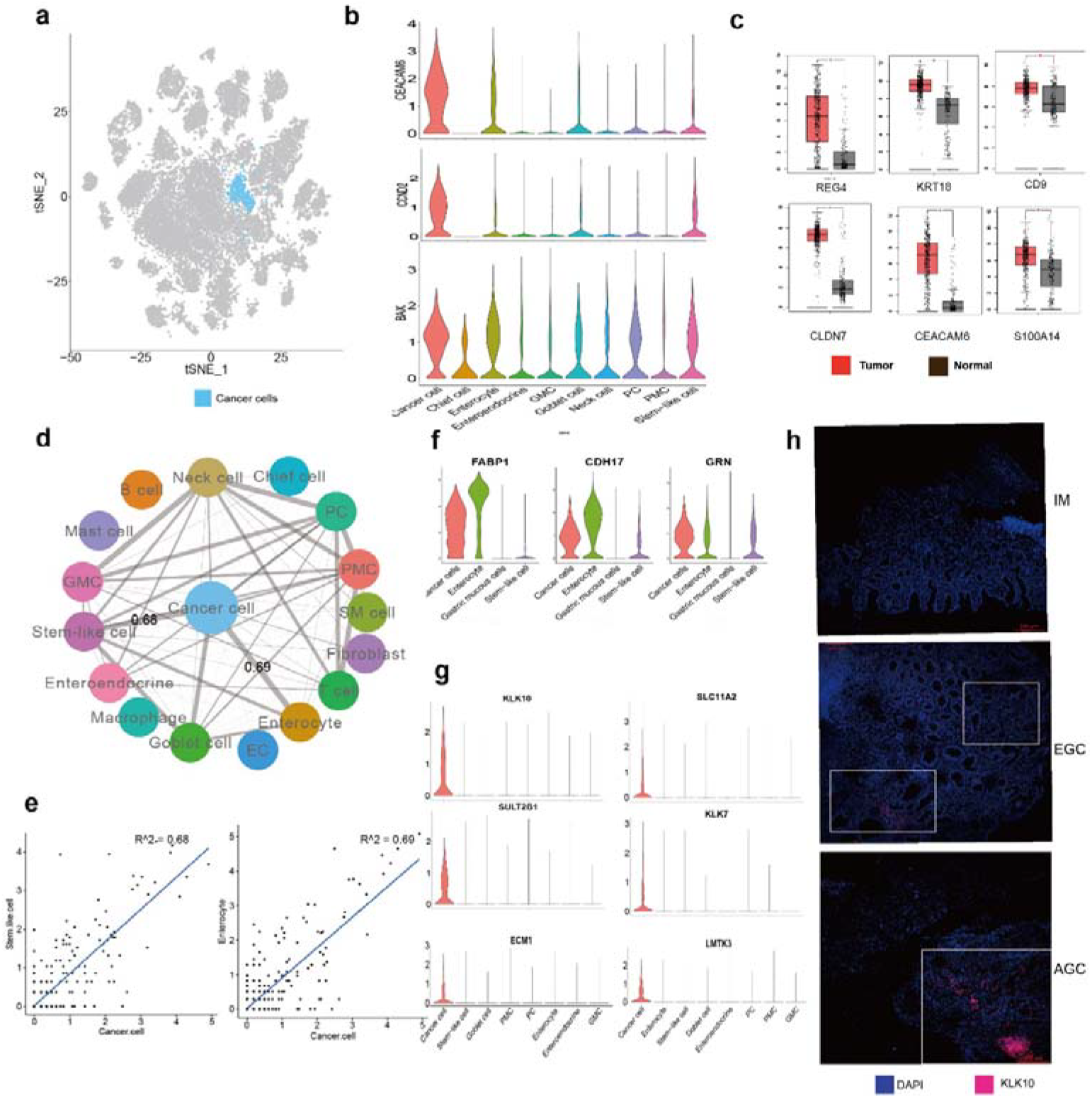
The scRNA profile of cancer cells in the EGC lesion. **a**. The t-SNE plot that showed the distribution of the ‘cancer cell’ cluster (blue, n = 798) in the atlas. **b**. Boxplot for the distribution of expression of the gastrointestinal cancer marker CEACAM6, cell cycle-related gene CCND2 and apoptosis-related gene BAX in diverse epithelial cell types, Mann–Whitney U-test, P∟< ∟1E-16. **c**. Boxplot of the differential expression for the putative cancer-cell-related top 6 upregulated genes in the gastric cancer datasets in TCGA. * FDR < 1e-5. **d**. The distance network among cell types in our dataset. The thickness of edges in the network was denoted as the pearson correlation coefficient between the centroids of any pair of cell types. **e**. The correlations between cancer cells and stem-like cells. **f.** Violin plots display the distribution of expression of previously reported EGC-related markers (FABP1 (left), CDH17 (middle) and GRN (right)) in cancer cells, enterocytes, gastric mucous cells and stem-like cells. **g**. The signature of EGC cells, Mann–Whitney U-test, P∟< ∟1E-16. **h**. The IF staining for the representative gene in the EGC signature, KLK10, in sections derived from IM, EGC, and advanced gastric cancer (AGC) biopsies.

Firstly, we confirmed the specific up-regulation of the gastrointestinal cancer marker gene, CEACAM6, and cell proliferation-related genes, BAX and CCND2, in the cancer cell clusters in the whole atlas (Figure 6b). Moreover, we identified an extended list of marker genes for the cell cluster (p < 0.01, fold change >1.5, Supplementary table 5) and KEGG enrichment analysis indicated that these marker genes were significantly enriched for cancer-related terms, including p53 signaling pathway (Supplementary table 9). We also confirmed the dysregulation of these identified marker genes in TCGA and other GC-related transcriptomic datasets, where most of them (60/80, Supplementary table 10) showed consistently significant up-regulation in malignant tissues in comparison to adjacent non-malignant ones (Figure 6c). Finally, we found that the cancer cluster was uniquely present in the EGC biopsy and rarely present in the two non-neoplastic biopsies from the same patient (Supplementary figure 15). These results re-confirmed cells in the cluster as the EGC cells.

Next, we focused on the transcriptional correlations between the cancer cell cluster and other cell populations. By representing each cell population by its centroid, we calculated the correlation scores for any pair of two cell clusters and visualized them in a network (Figure 6d). It revealed that the neoplastic cell cluster showed the greatest transcriptional similarity with the stem-like cell cluster and the enterocyte cluster (Figure 6e), which was consistent with the hallmarks for the intestinal type of gastric cancer.

In clinical practice, premalignant metaplastic sites usually reside in the surrounding of neoplastic mucosa, which confounds the precise identification of cancer cell-related gene profiles due to the transcriptional similarity between cancer cells and enterocytes. Actually, we found that some genes previously reported as GC-related early diagnosis markers, such as FABP1, CEACAM5 and CDH17, showed extensive expression in enterocytes or other cell types, suggesting their non-specific expression in cancer cells (Figure 6f). Therefore, based on the single-cell atlas, we defined a panel of high-confidence EGC-related marker genes by selecting those genes that showed significantly upregulated in cancer cells, while minimally expressed in other cell types (Methods). As a result, this panel consisted of seven genes, of which SLC11A2, KLK7 and KRT16 has not been reported to be involved in gastric tumourigenesis (Figure 6g). As a case, we validated the expression of the most up-regulated gene in the panel, KLK10, by immunofluorescence staining of human IM, EGC and advanced gastric cancer (AGC) biopsies, respectively. It revealed that the immunofluorescence staining of KLK10 showed negative in the IM sample, moderate in the EGC sample and intensely expressed in AGC sample. Thus, it suggested that these genes could be used as cancer cell-specific molecular markers and precisely recognizing the onset of GC cell development at early stages in clinical practice.

## Discussion

Although scRNA-seq profiles of other intact gastrointestinal organs, including the oesophagus and colon, have been addressed ^51,52^, those of the gastric mucosa, particularly mucosae in premalignant and malignant lesions, have not been demonstrated before. To our knowledge, this is the first study to define, in detail, a single-cell atlas of the gastric mucosae of patients with NAG, CAG, IM and EGC. For each lesion, we identified diverse cell types, and defined core gene expression signatures for these cell types. We also analysed the transcriptomic changes of some cell types, including mucous cells and enteroendocrine cells, across different lesions. Additionally, goblet cells and neoplastic cells, the representative cellular characteristics for the onset of IM and GC, were analysed in depth for identifying cell-type-specific markers that are potentially applicable in clinical practice.

Although IM has been demonstrated as the precursor of GC, the link between metaplastic lineage cells and the evolution of intestinal-type GC has remained unclear ^53–57^. In our study, we observed that both EGC and IM biopsies contained cells expressing the stem cell-related gene OLFM4, whereas CAG biopsies did not. OLFM4 has been demonstrated as a robust stem cell marker in the human intestine and marks a subset of colorectal cancer cells ^29^. Thus, this indicates that cancer cell lineages might contain similar progenitor cells as those of IM cell lineages, other than gastric cell lineages. Further studies with single-cell DNA sequencing techniques could help confirm the relationship between metaplastic and neoplastic cell lineages, which might facilitate our understanding of the mechanism underlying gastritis-induced GC.

Early detection of GC is crucial in clinical practice because the early diagnosis of gastric cancer in patients leads to a significant improvement in prognosis compared to that of those diagnosed at the advanced stage of GC patients ^58–60^. However, many of well-documented GC-associated markers, such as carcinoembryonic antigen (CEA) and carbohydrate antigen 19-9 (CA19-9), lack sufficient sensitivity and specificity to enable perform early detection ^58,61^. Here, we identified the ‘neoplastic cell cluster’ based on single-cell transcript expression from multiple biopsies, including neoplastic and non-neoplastic biopsies, from the same patient with EGC. We observed that the neoplastic biopsy comprised multiple cell lineages besides neoplastic cells, and the identified neoplastic cells showed relatively similar transcriptional patterns to those of other cell types, such as enterocytes and gastric pit cells. Thus, the use of bulk sample-based sequencing technology would confound the precise identification of diagnostic markers for EGC. Therefore, we speculated that single-cell sequencing technology might provide a novel and high-resolution method for the identification of molecular markers for use in precisely detecting early neoplastic sites and the onset of rare neoplastic cells in the clinic.

There are also some limitations in our study. First, the spatial origin of individual cells along the gastric gland axis is lost when the tissue is dissociated for scRNA-seq sequencing. This issue could be addressed by combining laser-capture-microdissected technology and scRNA-seq to dissect epithelial cells in different regions along the gland axis^62^. Second, although multiple biopsies were collected from distinct sites in the same patient to mimic different lesions in the cascade from CAG to GC, longitude data based on monitoring a patient during the development from premalignant to malignant lesions might more precisely reflect the process underlying inflammation-induced tumourigenesis.

In summary, we constructed a single-cell transcriptome atlas of gastric premalignant and early-malignant mocusa for the first time. With the atlas, we characterized the expression patterns of diverse cell types in each lesion and analysed their changes across lesions. Of note, we also identified a panel of early neoplastic cell-specific marker genes, providing a molecular basis to precisely diagnose EGC. Together, our findings provide unparalleled insight into the cellular heterogeneity of gastric mucosae with different types of premalignant lesions and malignant lesions, which might be helpful for identifying markers for cancer prevention and facilitate our understanding of GC pathogenesis.

## Materials and Methods

### Sample collection and processing

In this study, gastric mucosae and clinical information including traditional Chinese medicine information were collected from patients diagnosed with gastritis, intestinal metaplasia and early gastric cancer, after written informed consent was obtained, from China-Japan Friendship Hospital (CJFH), China. The protocol in our study was approved by The Ethical Committee of China-Japan Friendship Hospital. For each patient, apart from tissue biopsies for conventional histological examination, additional one or more biopsies were taken from the gastric antrum for correlative studies. For the early gastric cancer patient, mucosa biopsies were surgically removed from three sites, with one at the cancer site and the remaining two at remote sites with different severity in IM; For other patients, biopsies were all obtained from one or two sites by conventional upper gastrointestinal endoscopy performed by specialist endoscopists using standard forward viewing videoscopes. All of the fresh resected biopsies were divided into two equal parts, of which one was processed for the single-cell sequencing experiment and another was for other experiments, including the pathological grading, immunohistochemical studies. Each biopsy sample was assessed by two independent experienced pathologists for the degree of chronic gastritis, the presence of gastric atrophy (GA), and intestinal metaplasia (IM). IMs were classified as mild or severe, based on hematoxylin and eosin (H&E) staining and MUC2-based immunofluorescence staining. Because IMs are heterogeneous, classifications were based on the predominant type of IM for each biopsy. Cancer in the T0 stage (carcinoma in situ) was defined as the early gastric cancer (EGC) in this study according to the 7th edition of the American Joint Committee on Cancer (AJCC) TNM stage system. The presence of Hp infection was determined on the basis of the consensus result from histology analysis and the C13 urea breath testing (UBT). The endoscopic image of each site in the stomach and the tongue image of each patient were also collected, as shown in Supplementary figure 3.

All of the fresh resected mucosae were processed as described below. Briefly, each biopsy was minced with the Iris scissor into small pieces and digested for 30min at 37°C, 800 rpm, with a digestion solution containing PBS and collagenas II and IV (1.5mg/ml, gibco). The cell suspension was further filtered through 45-um nylon mesh to remove cell aggregates and re-suspended in PBS with 10% FBS. Cell sorting was performed with a MoFlo XDP (Beckman coulter). Dead cells were eliminated by excluding Sytox-positive (Membrane Permeability Dead Cell Apoptosis Kit with PO-PRO-1 and 7-Aminoactinomycin D, Life Technologies) cells, which increased the efficiency of sorting robust, live cells for single-cell experiments.

### Library preparation and sequencing

Library preparation was performed according to instruction in the 10× Chromium single-cell kit. The libraries were then pooled and sequenced across six lanes on a HiSeq4000.

### Quality control and RNA-seq data pre-processing

We firstly processed 10X genomics raw data by the Cell Ranger Single-Cell Software Suite (release 2.0), including using *cellranger mkfastq* to demultiplexes raw base call files into FASTQ files and then using *cellranger count* to preform alignment, filtering, barcode counting, and UMI counting. The reads were aligned to the hg19 reference genome using a pre-built annotation package download from the 10X Genomics website. The output from different lanes was final aggregated using ‘cellranger aggr’ with default parameter setting.

Then we mapped UMIs to genes, followed by removing low-quality cells. Cells would be flagged as poor-quality ones if they met one of the following thresholds: 1) the number of expressed genes lower than 400 or larger than 7000; 2) 20% or more of UMIs were mapped to mitochondrial or ribosomal genes. Cells meeting the latter threshold were usually non-viable or apoptotic. According to the previous study, we also excluded cells with barcodes appeared in more than one sample with respect to index swapping. As a result. We detected 22,882 genes in a total of 31,164 cells, as shown in Supplementary table 3.

Then, we utilized functions in the Seurat package^63^ to normalize and scale the single-cell gene expression data. It was firstly normalized by “NormalizeData” function with setting normalization method as “LogNormalize”. In detail, the expression of each gene i in cell j was determined by the UMI count of gene i divided by the total number of UMI of the cell j, followed by multiplying 10000 for the normalization and the log-transformed counts were then computed with base as 2. We then removed the uninteresting sources of variation by regressing out cell-cell variation within gene expression driven by batch, the number of detected UMI, mitochondrial gene expression, as well as ribosomal gene expression, which was implemented by “ScaleData” function. Finally, the corrected expression matrix was used as an input for further analysis.

### Dimension reduction & Cell clustering & Annotation

We then restricted the corrected expression matrix to the subsets of highly variable genes (HVGs), and then centered and scaled values before performing dimension reduction and clustering on them. Methodologically, the highly variable genes (HVGs) in single-cell data were selected by first fitting a generalized linear model to the mean-dependent trend to the gene-specific variance to all genes, and selecting genes that deviated significantly from the fitted curve. It was implemented by “FindVariableGenes” function in the Seurat package by setting the valid value of average expression as a range from 0.05 to 5 and that of dispersion as no less than 0.5. It left 1,258 genes as HVGs. We then used the “RunPCA” function in the Seurat package to perform the principle component analysis (PCA) on the single-cell expression matrix with genes restricted to HVGs. Given that many principle components explain very low proportion of the variance, the signal-to-noise ratio can be improved substantially by selecting a subset of significant principle components. The number of significant principal components were determined by using the permutation test, implemented by permutationPA function from the jackstraw R package (https://cran.r-project.org/web/packages/jackstraw). The analysis identified 50 significant principal components as a result and scores from only these principle components were used for further analysis.

We then utilized the “FindClusters” function in the Seurat package to conduct the cell clustering analysis through embedding cells into a graph structure in PCA space. Due to the large number of cells in our study, we set the parameter resolution as 2. This identified a total of 35 clusters. We then performed a post-hoc test by merging clusters with less than 10 differentially expressed genes (p < 0.01 & fold change > 2). We annotated cell clusters based on the expression of curated known cell markers, as shown in the Supplementary table 4. Although some clusters were identified as isolated ones, we found they consistently expressed the same cell marker. Isolated clusters C25, C6, C31 and C23 were consistently expressed the gland mucous cell (GMC) marker MUC6 and thus we viewed all of them as the GMC cluster; In addition, isolated clusters C22, C26, C24, C0 were consistently express the gastric pit mucous cell (PMC) marker MUC5AC and thus we viewed all of them as the PMC cluster. The t-SNE embedding was computed using the ‘TSNEPlot’ function with default settings.

When processing the specific cell lineages including gastric mucous-secreting cells, enteroendocrine cells, goblet cells and cancer cells, we re-run the above pipeline with keeping the same parameter setting in the “FindVariableGenes” function and choosing the top 20 significant principal components as the features in the PCA space.

In addition, we annotated enteroendocrine cell clusters with the dominate expression pattern of canonical endocrine cell markers. Briefly, we first mapped the expression of canonical endocine markers to each putative cluster and count the proportion of marker-expressing cells for each cell type in the cluster. Then, we removed the low-abundance cell types with lower than 20% and termed the cluster with combining the name of remaining cell types.

### Differential expression analysis

Differential gene expression analysis was performed using the ‘FindMarkers’ function, which performs differential expression based on the non-parameteric Wilcox rank sum test for two annotated cell groups. The marker genes visualized in the Figure 1d were identified by using the ‘FindAllMarkers’ function in Seurat with settings on genes with at least 2-fold up-regulation, comparing to the remaining cells.

### Gene set enrichment analysis

We conducted the gene set enrichment analysis for concerned gene list by the tool Enrichr^64^, with which the enriched Kyoto Encyclopedia of Genes and Genomes (KEGG) pathways and Gene Ontology (GO) terms were derived.

### Inferring high-risk genes for gastritis and gastric cancer

We inferred the high-risk genes for gastritis or gastric cancer by utilizing the bioinformatics algorithm, CIPHER^34^, which prioritizes disease-related genes on a genome-wide scale according to network correlations between disease phenotypes and corresponding genes. Taking disease terms (gastritis: MIM137280 and gastric cancer: MIM137280) as inputs, we selected genes in the top 1000 candidates in prediction profiles as gastritis and gastric cancer-related genes, respectively, followed by mapping them to standard gene symbols.

### Construction of single-cell transcriptome network

We constructed the single-cell transcriptome network underlying gastric premalignant and early gastric cancer by connecting signature genes with known protein-protein interactions (PPIs) documented in STRING database (version 10, http://string-db.org)^65^ for each lesion. Herein, the signature genes for each lesion were derived by merging up-regulated genes in cell types present in the lesion. As a result, we colored each signature gene according to cell types where it preferentially expressed. Isolated network nodes were removed from the result.

### Immunofluorescence staining

Formalin-fixed, paraffin-embedded Sections (4∟μm) were deparaffinized in xylene and then hydrated in graded alcohol. EDTA (pH 8.0) was used for antigen retrieval in boiling water. The specimens were blocked by 3% H2O2 for 30min and then by 3% Albumin BovineV (Solarbio) for 30min. Samples were incubated with primary antibody overnight at 4°C, rinsed in PBS, then detected by fluorescent secondary antibodies (anti-mouse IgG;1:200;abcam) for 30 min at 37°C,rinsed in PBS, and finally stained with DAPI for 10min. Fluorescence microscopy was performed using a Zeiss 780 confocal microscope.

### Data availability

The single-cell RNA sequencing data have been deposited in the Gene Expression Omnibus (GEO) database with the accession code as GSEXXX. All computational analyses in this study were conducted in R (Version 3.4.3) using standard functions unless otherwise indicated in the paper. Codes are available online at http://bioinfo.au.tsinghua.edu.cn/member/pzhang/scstomach.r

## Acknowledgements

We wish to thank Jin Gu and Yang Chen for their comments and suggestions; Kui Hua, Dongfang Wang, Hua Chen and Qingyang Ding for their technical discussions.

## Author contributions

Peng Zhang designed most experiments, collected biopsies, analyzed data and wrote the manuscript. Mingran Yang designed experiments, collected biopsies, performed immunohistochemical staining and immunofluorescence experiments and contributed to the writing of the manuscript. Yiding Zhang performed data analysis, made figures and contributed to the writing of the manuscript. Shuai Xiao, Xinxing Lai, Feiran Zhang and Aidi Tan contributed to collecting biopsies. Shiyu Du contributed to collecting biopsies and designing experiments. Shao Li was responsible for the overall conception and design of the study as well as in drafting and revising the manuscript.

## Sources of Funding

This work was supported by grants from the National Natural Science Foundation of China (91729301, 81630103) and The Project of Tsinghua-Fuzhou Insititute for Data Technology (TFIDT2018001).

## Disclosures

None.

## Supplementary Information

**Supplementary Figure 1.**
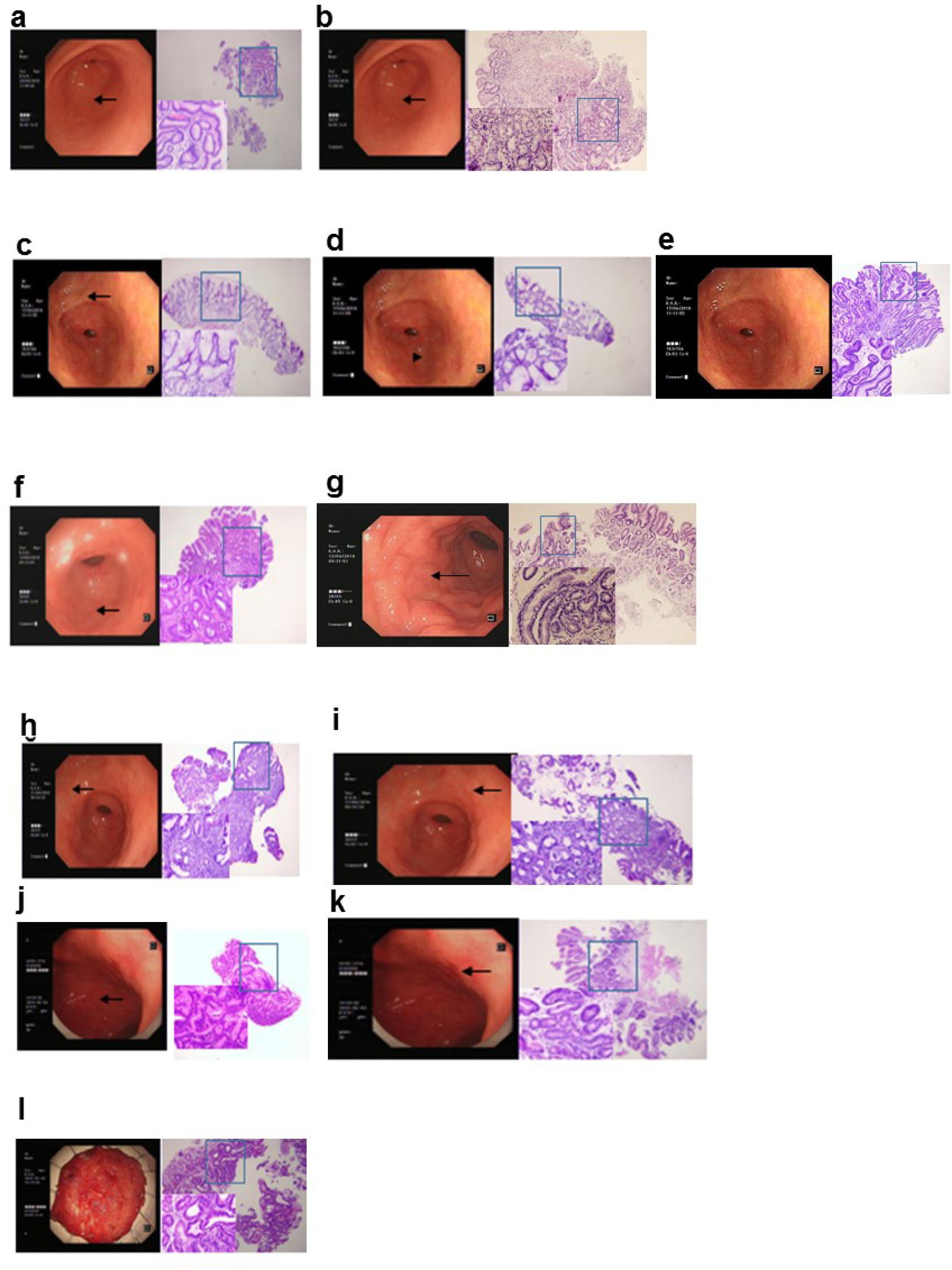
Gastroscope images and the hematoxylin and eosin (H&E) staining staining of samples in this study. Arrows show the sites of Gastroscopic biopsies. a, NAG1 (P1); b, NAG2 (P2); c, CAG1 (P3); d, CAG2 (P3); e, CAG3 (P4); f, IMW1 (P5); g, IMW2 (P6); h, IMS1 (P7); i, IMS2 (P7); j, IMS3 (P8); k, IMS4 (P8); l, EGC (P8)

**Supplementary Figure 2.**
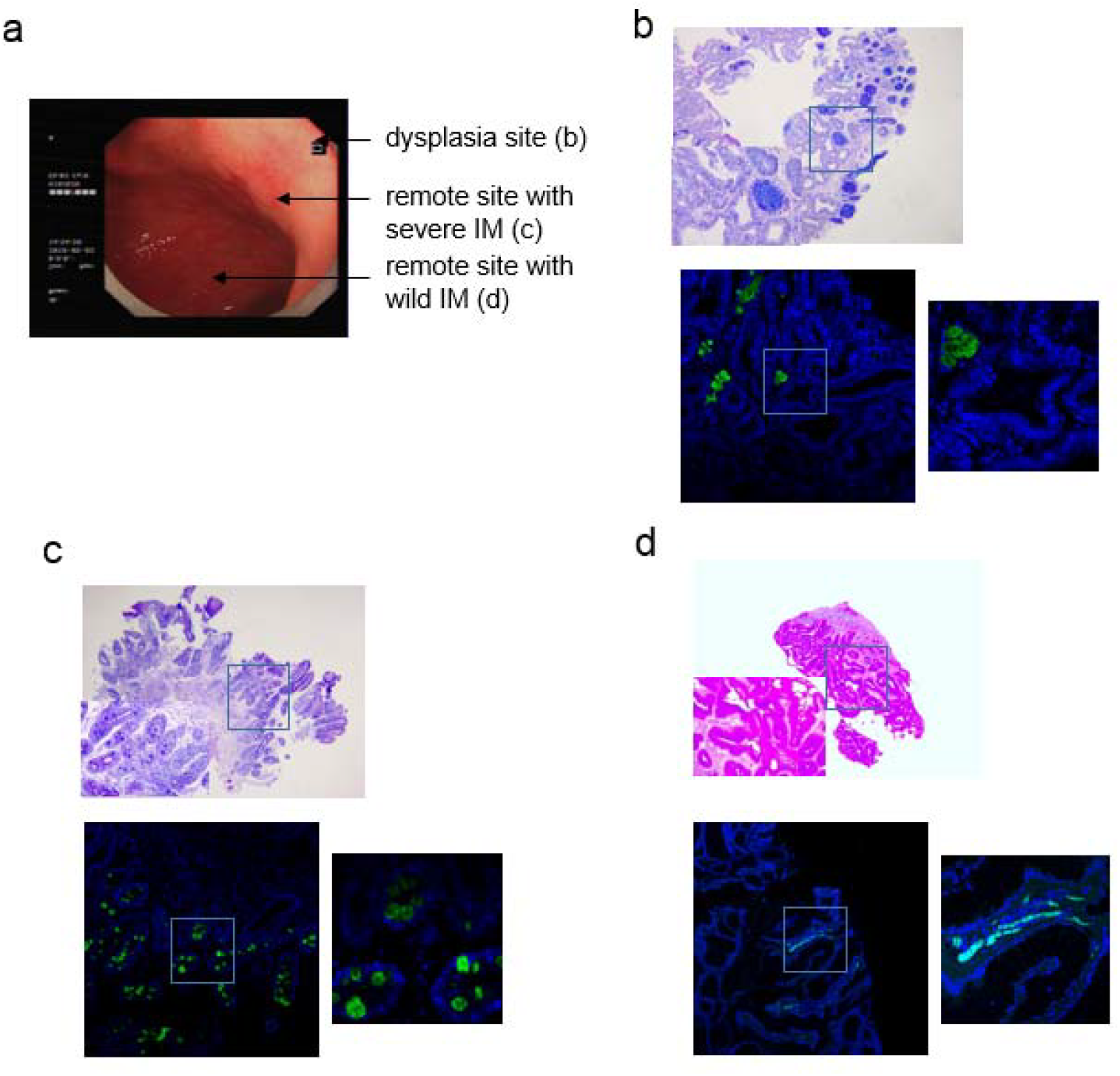
The the AB-PAS staining and MUC2-based immunofluorescence (IF) staining for the three biopsies collecting at three distinct sites from the early-malignant patient (P8). a, the three sites of biopsies in the gastric antrum of P8, including the neoplastic site, the remote site with severe IM and the remote site with wild IM. b-d, the AB-PAS staining (upper) and Immunofluorescence staining (bottom) of gastric tissue sections at the neoplastic site (b), remote site with severe IM (c) and remote site with wild site (d) (original magnification, 40x and 200x).

**Supplementary Figure 3.**
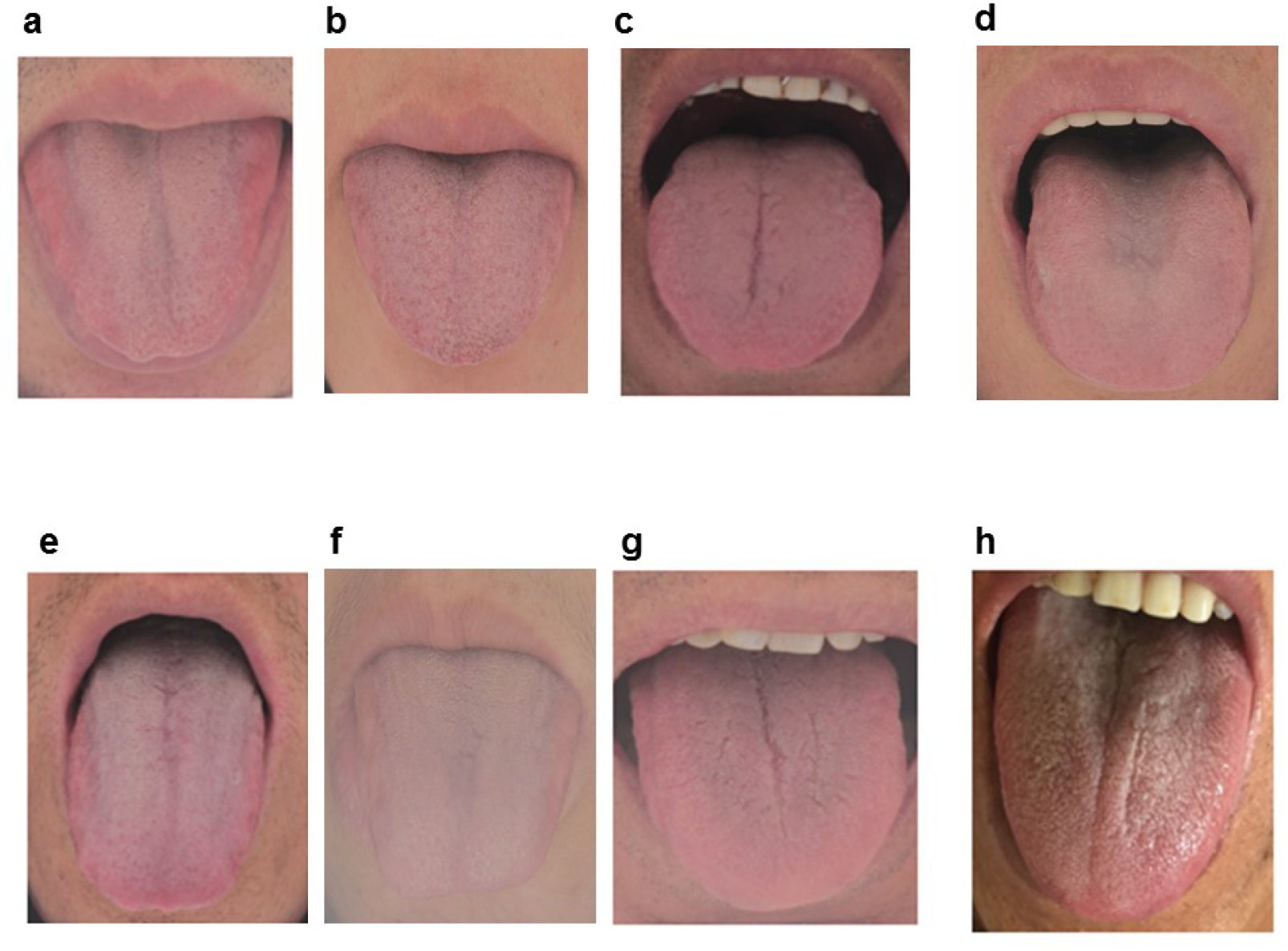
Tongue images of patients in the study. a, P1; b, P2; c, P3; d, P4; e, P5; f, P6; g, P7; h, P8.

**Supplementary Figure 4.**
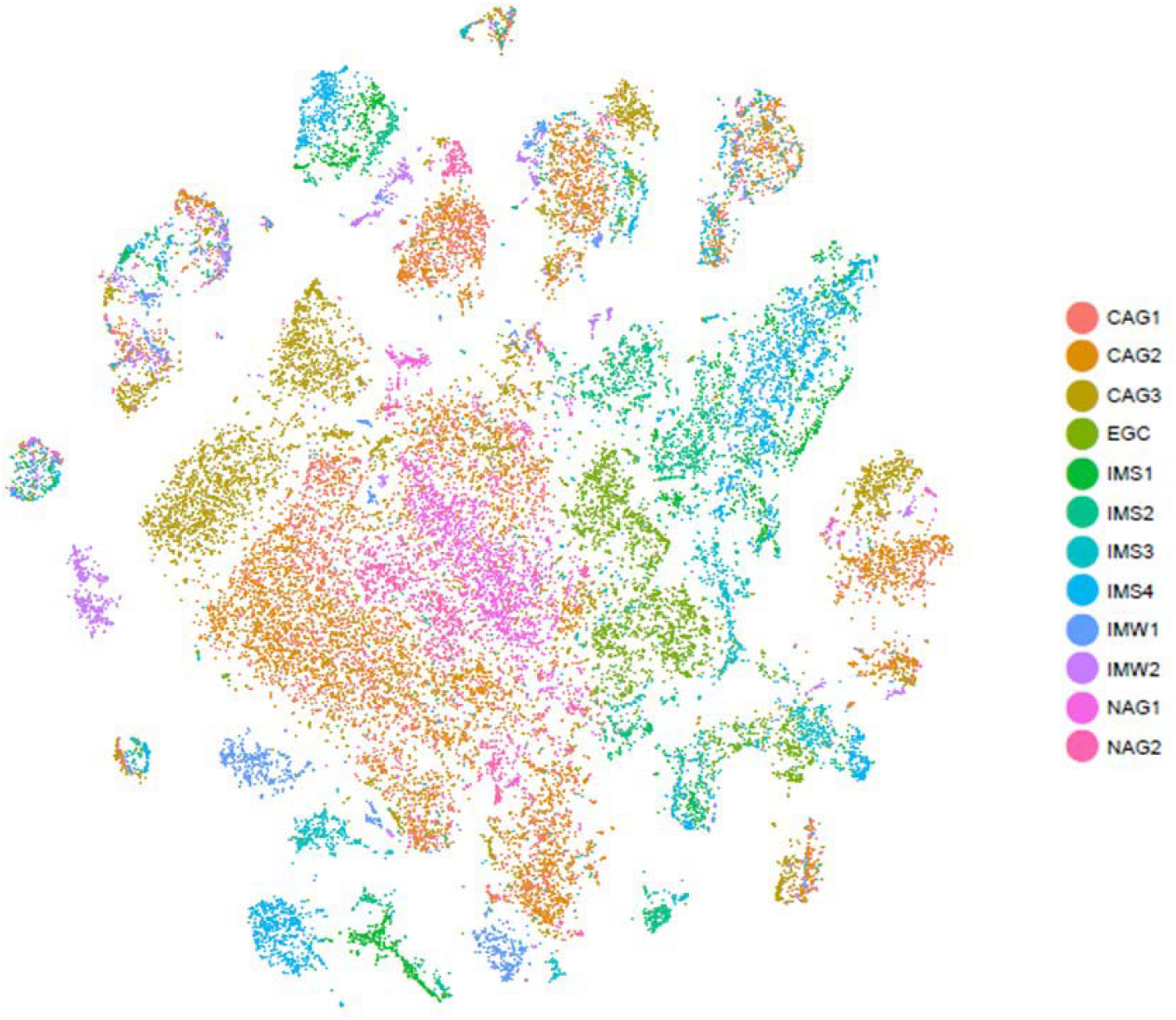
The t-SNE plot show the cell distribution among multiple batches.

**Supplementary Figure 5.**
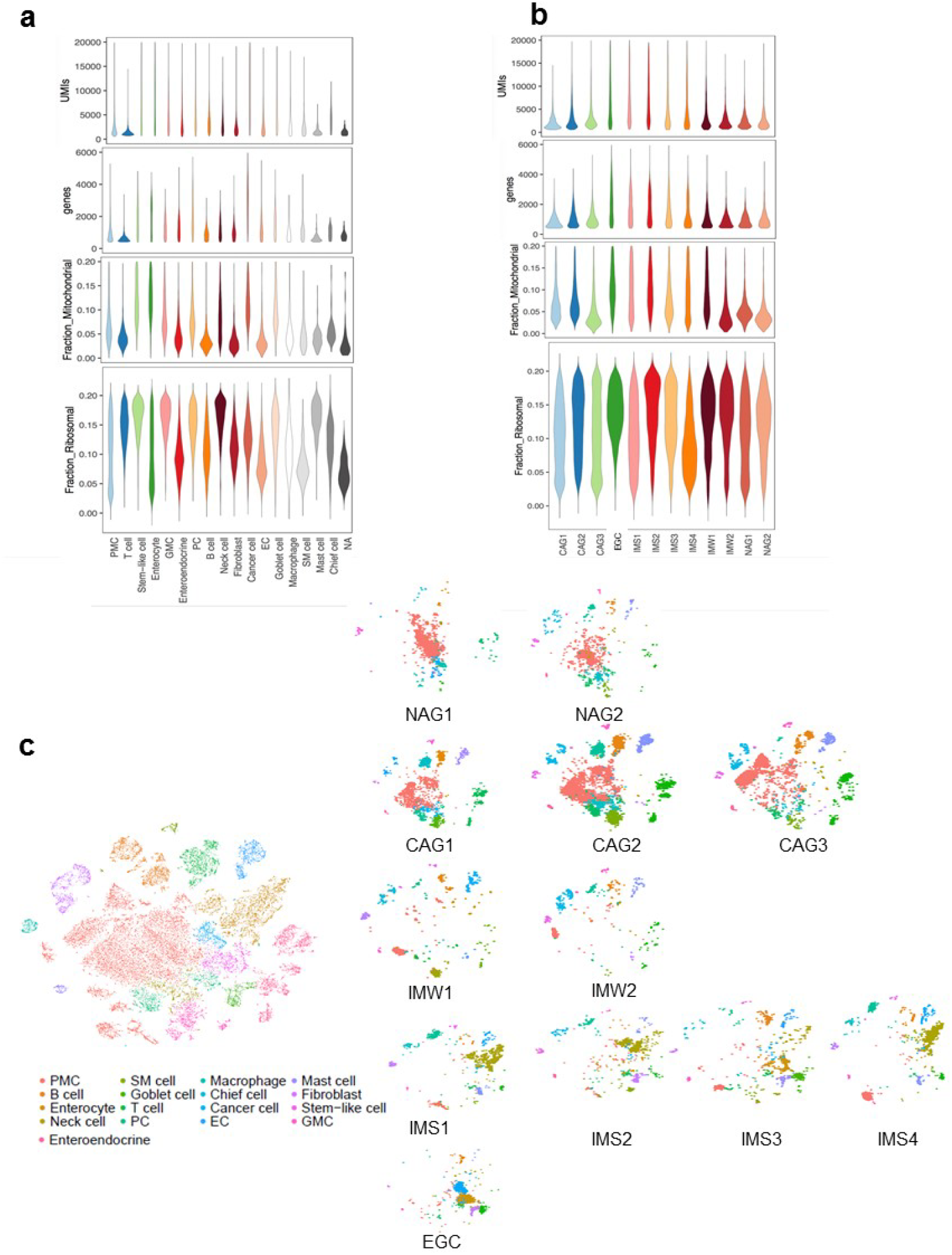
Consistency of cell capture and identification in biopsies from patients with different lesions. **a**, Number of unique molecular identifiers (nUMI) and genes identified, and fraction of reads mapping to mitochondrial or ribosomal genes across identified cell types. b, nUMI and genes identified, and fraction of reads mapping to mitochondrial or ribosomal genes across patient samples. c, t-SNE plot as in Fig. 1b coloured by cell types across all patients and then separated by sample.

**Supplementary Figure 6.**
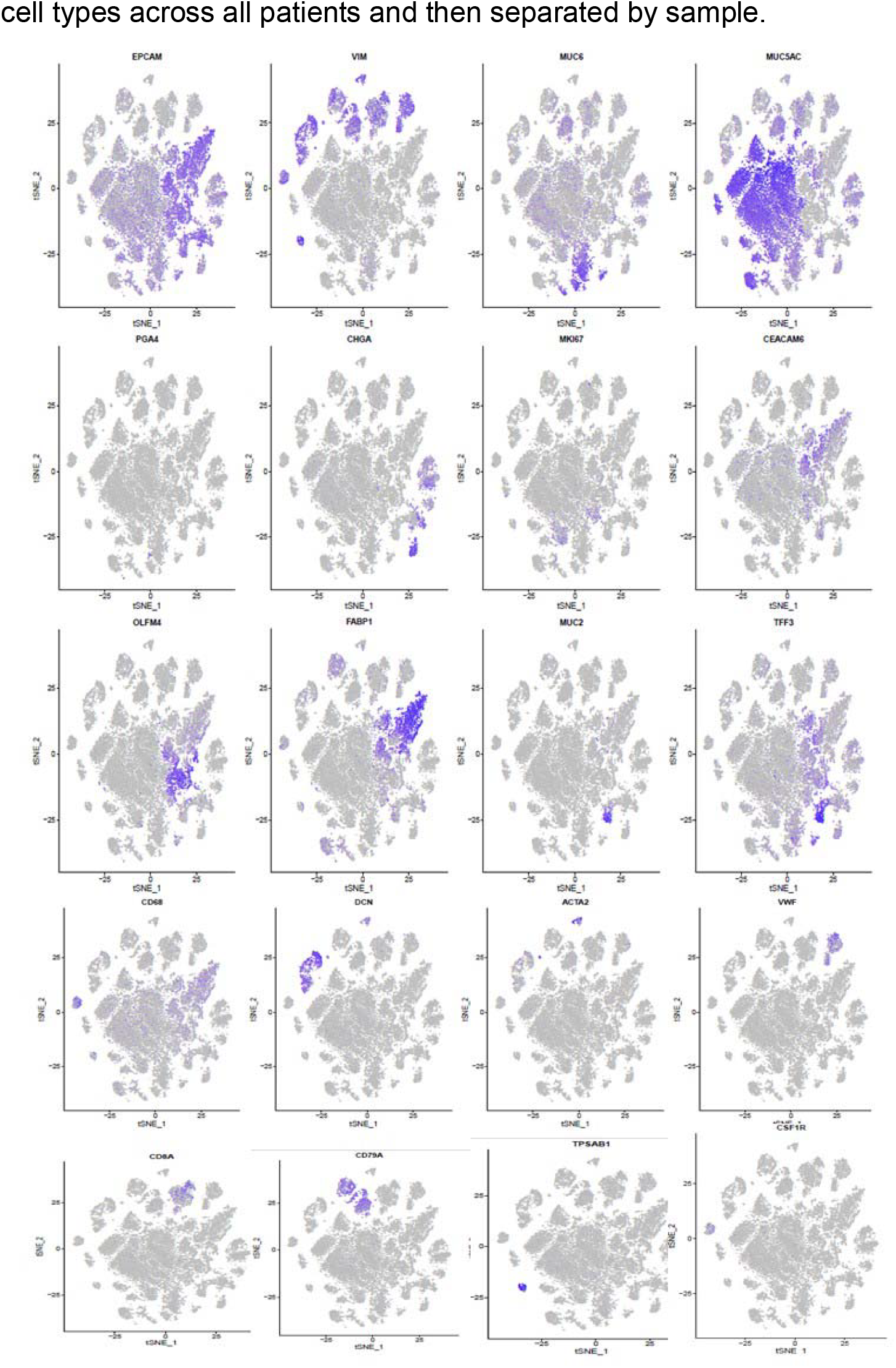
Smaller t-SNE plots show expression of known cell markers with cells colored according to the relative expression of the gene shown (blue = high, gray = low).

**Supplementary Figure 7.**
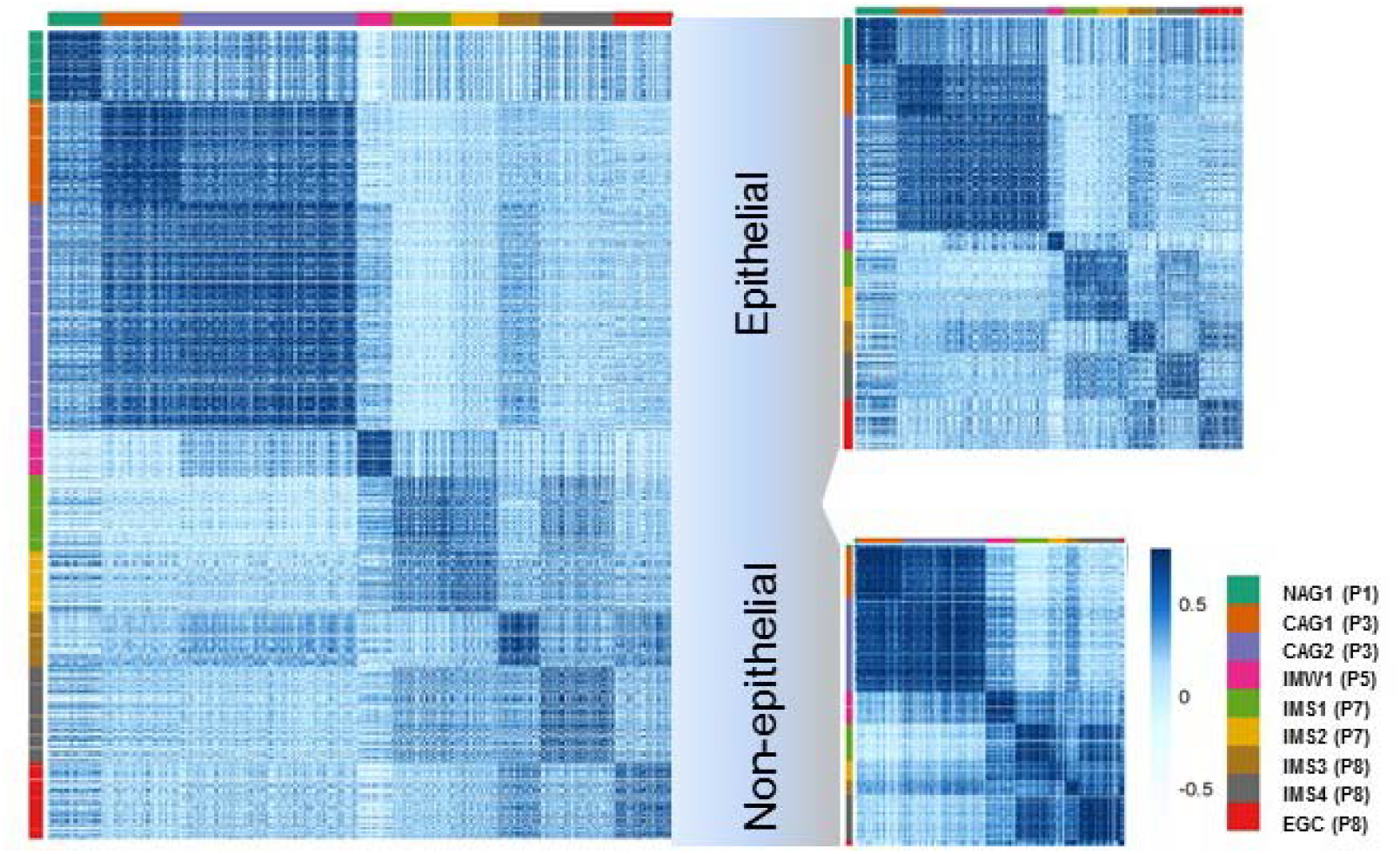
Cell-to-cell correlation matrixes for all single cells (left), epithelial cells (upper right) and non-epithelial cells (lower right). Each row and column represents single cells. Samples were represented in the color panel on the bottom right side.

**Supplementary Figure 8.**
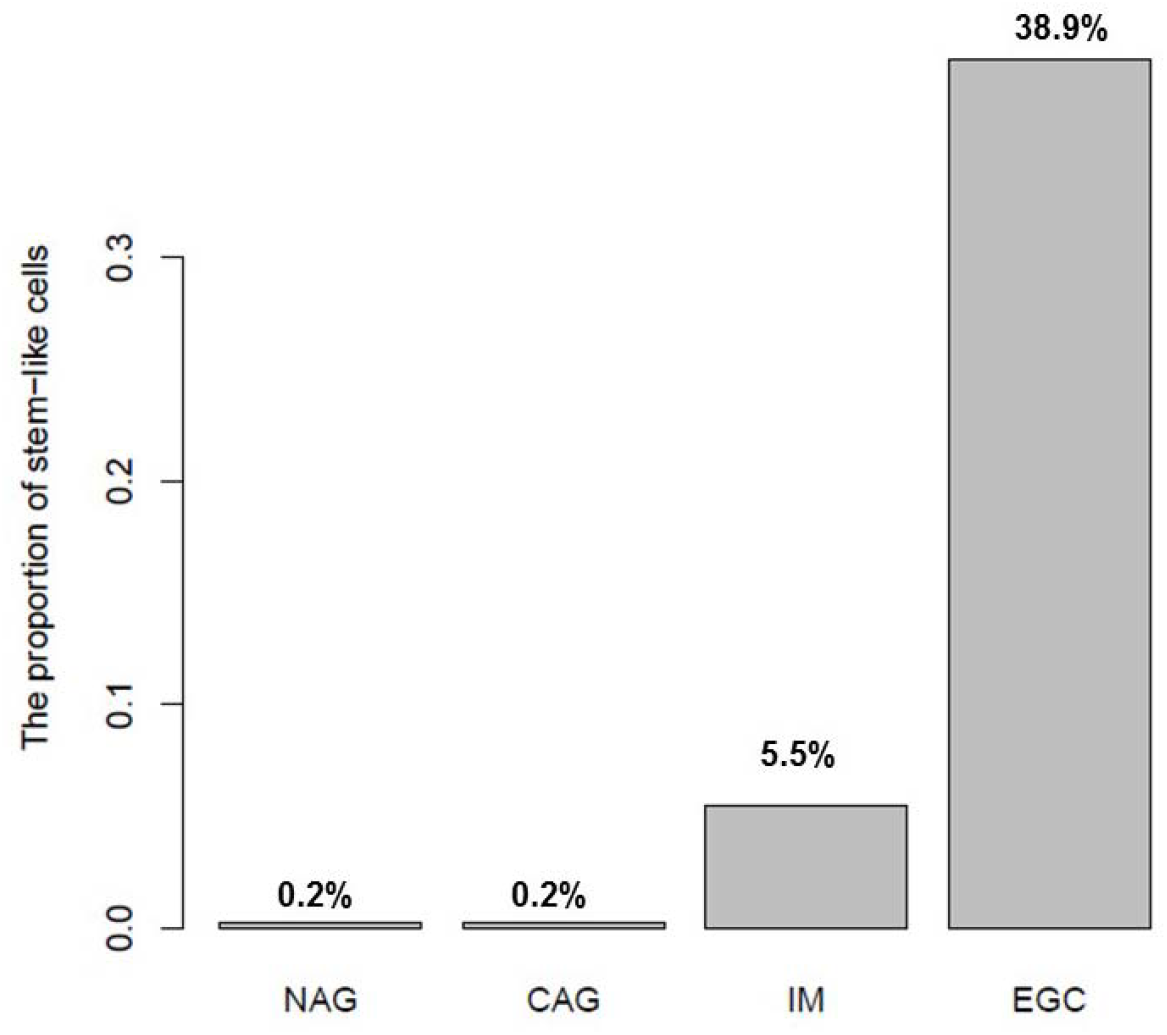
The proportion of stem-like cells in each lesion.

**Supplementary Figure 9.**
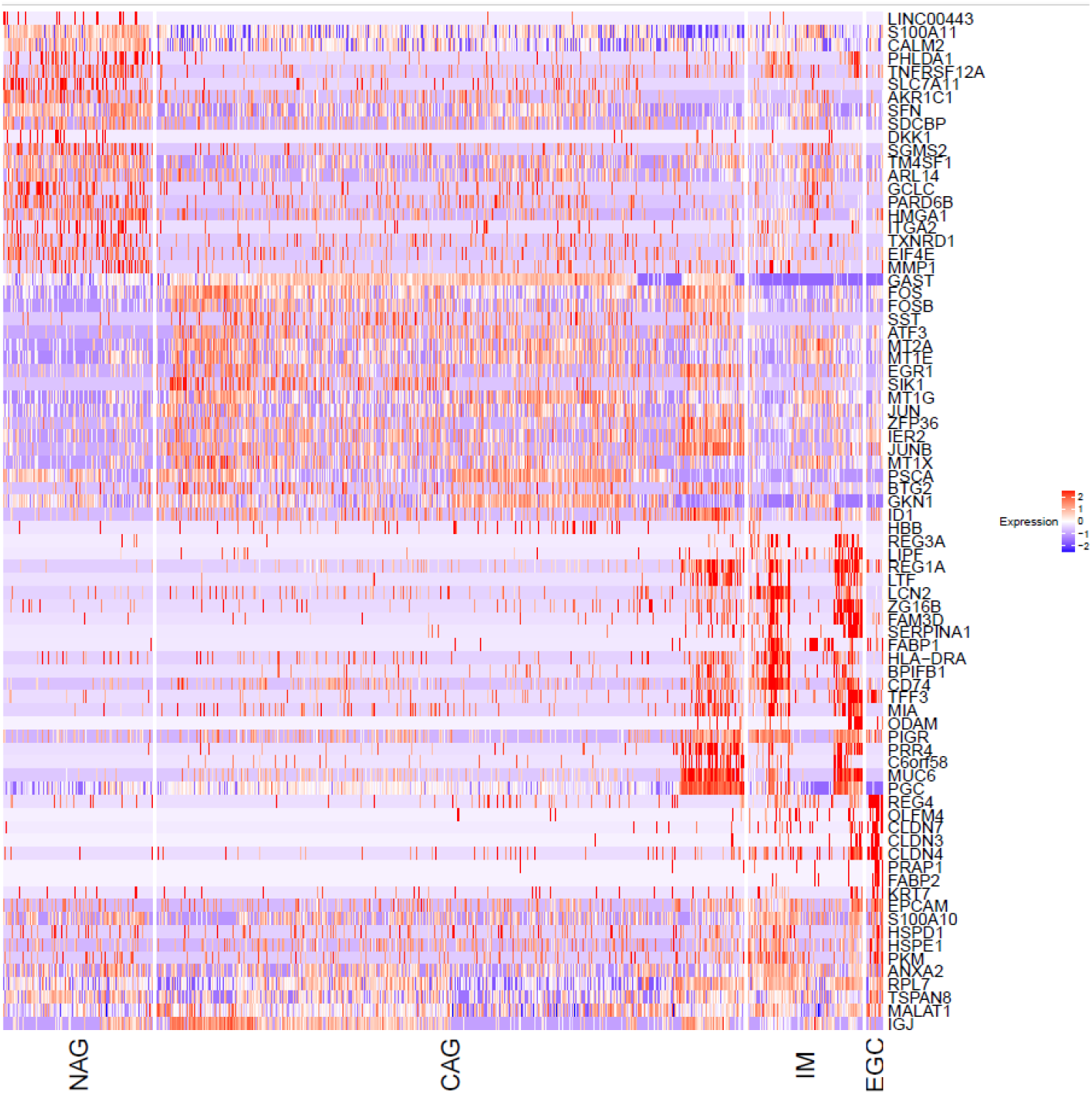
The heatmap for the expression profile of pit mucous cells (PMCs) cross different lesions.

**Supplementary Figure 10.**
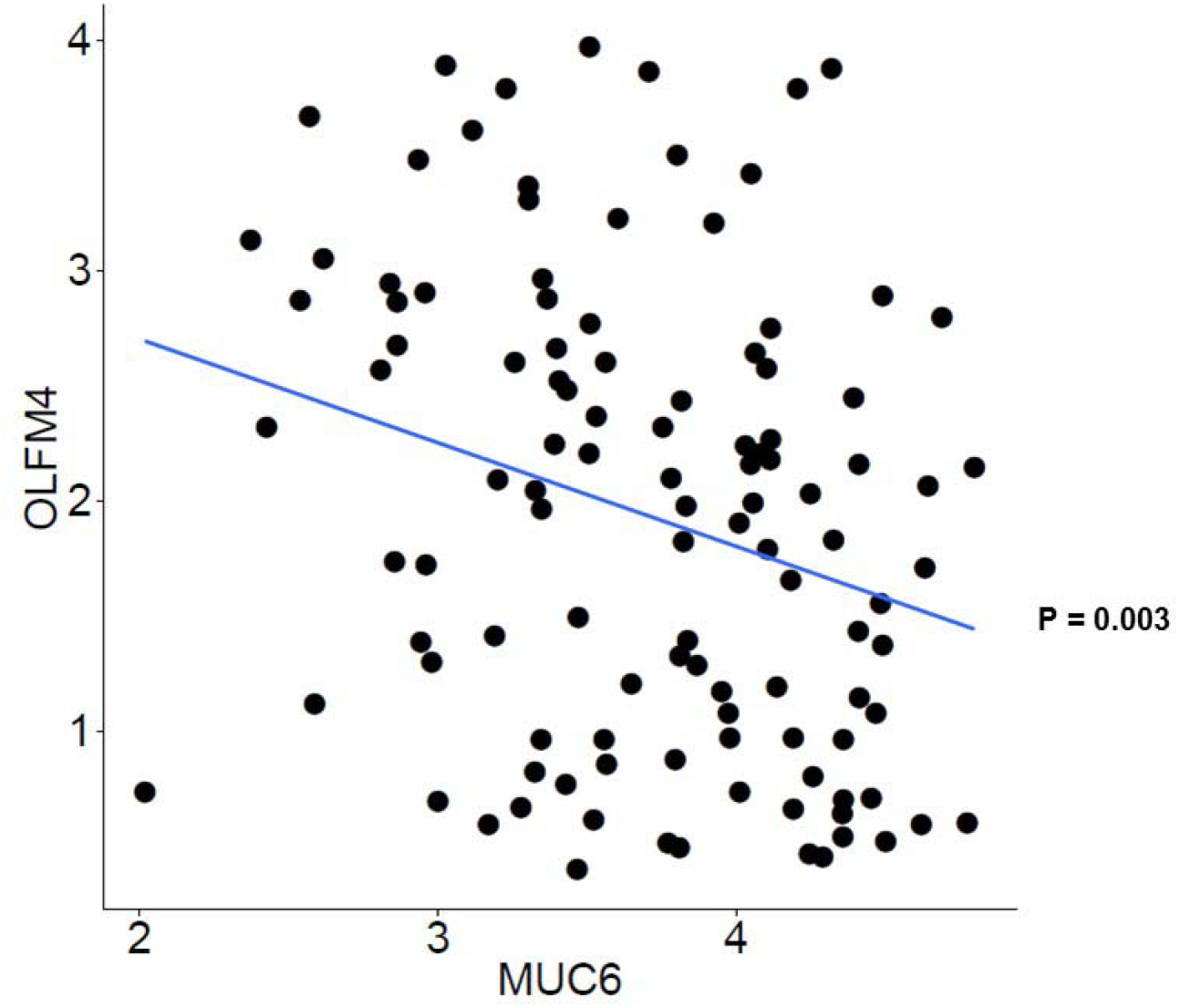
The co-expression of MUC6 and OLFM4 in individual GMCs in the IM lesion.

**Supplementary Figure 11.**
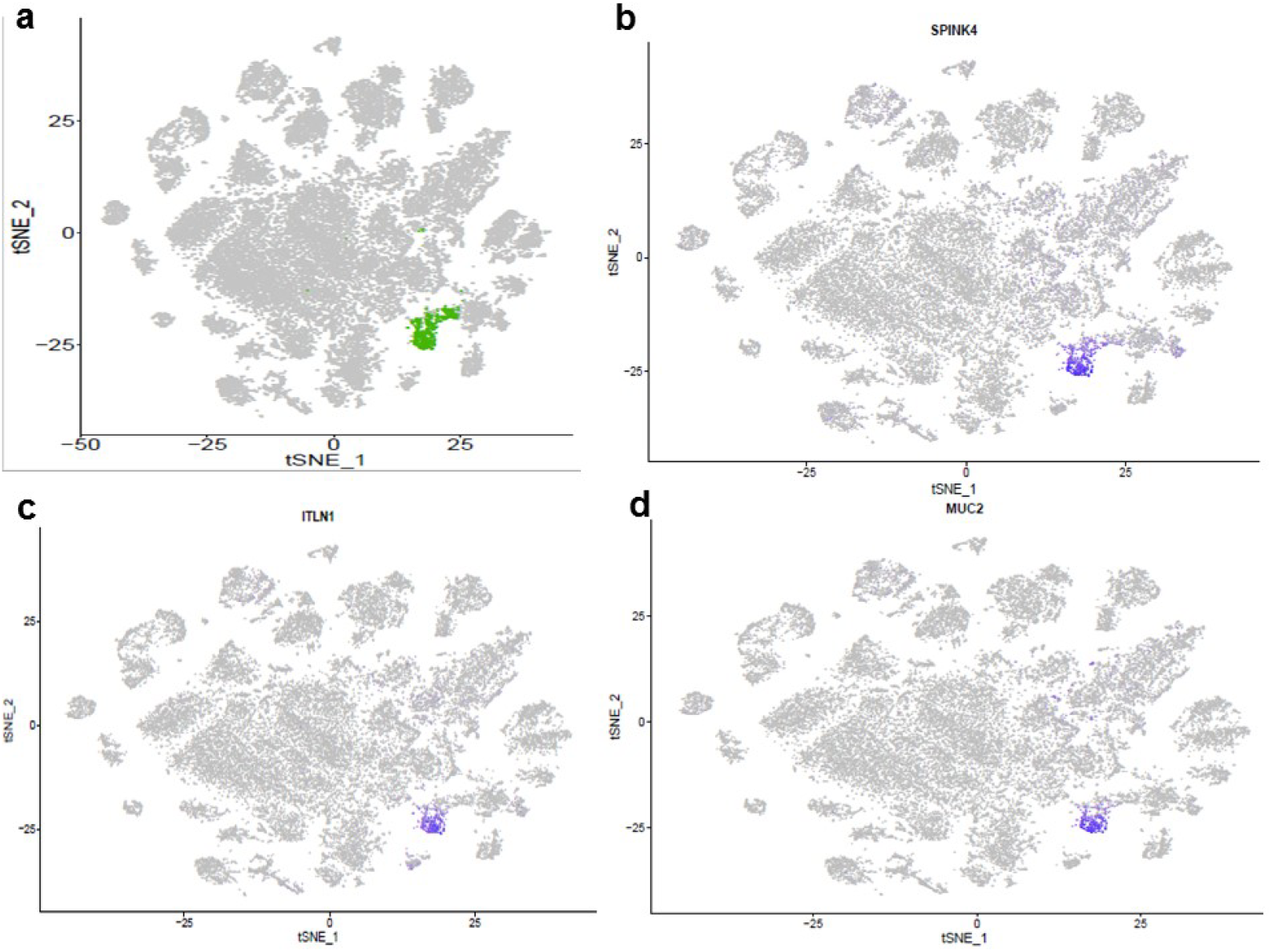
The t-SNE plot showing the distribution of ‘goblet cell’ cluster (a), with marked by the expression of SPINK4 (b), ITLN1(c) and MUC2 (d), respectively (blue = high, gray = low).

**Supplementary Figure 12.**
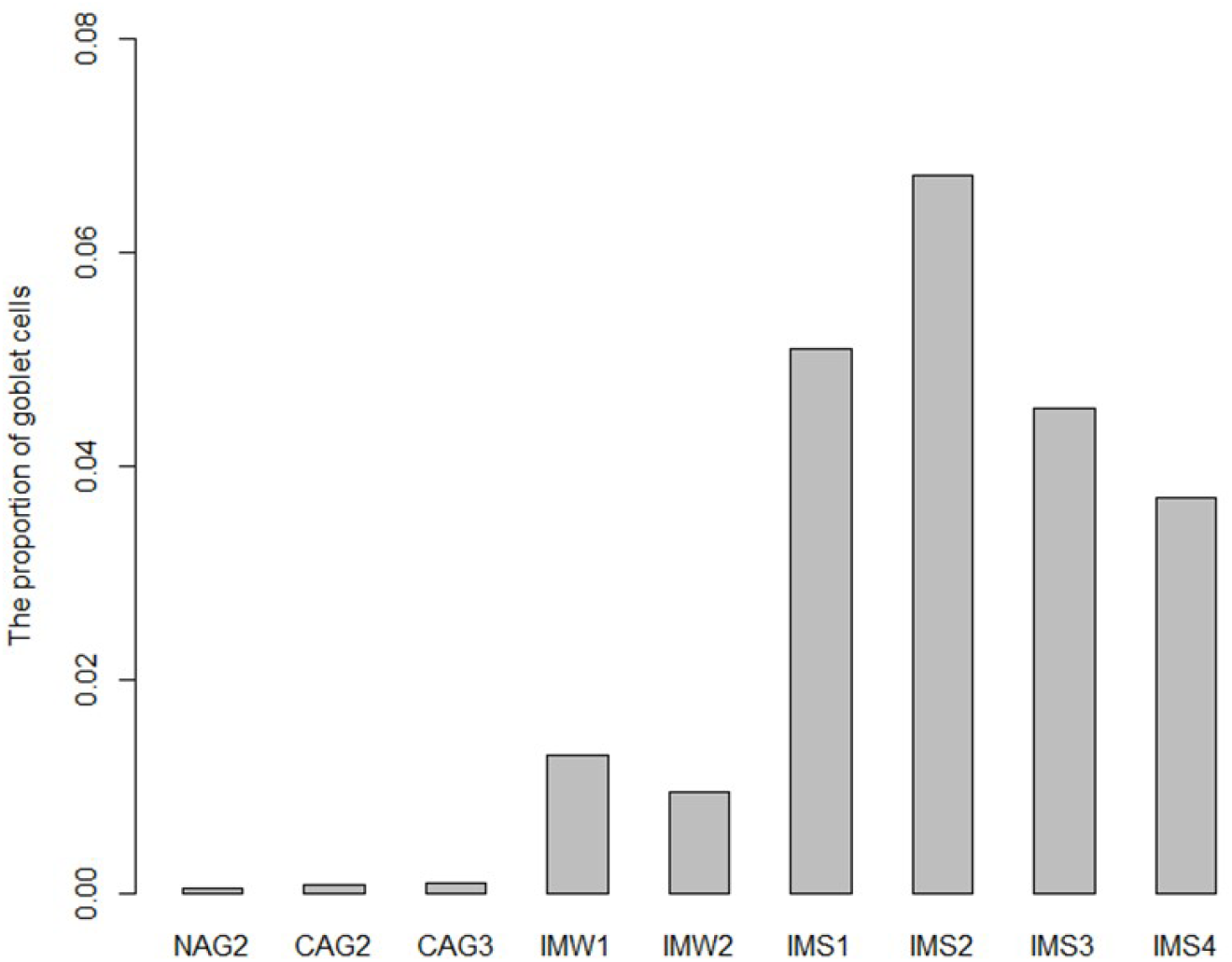
The proportion of goblet cells in each premalignant lesion.

**Supplementary Figure 13.**
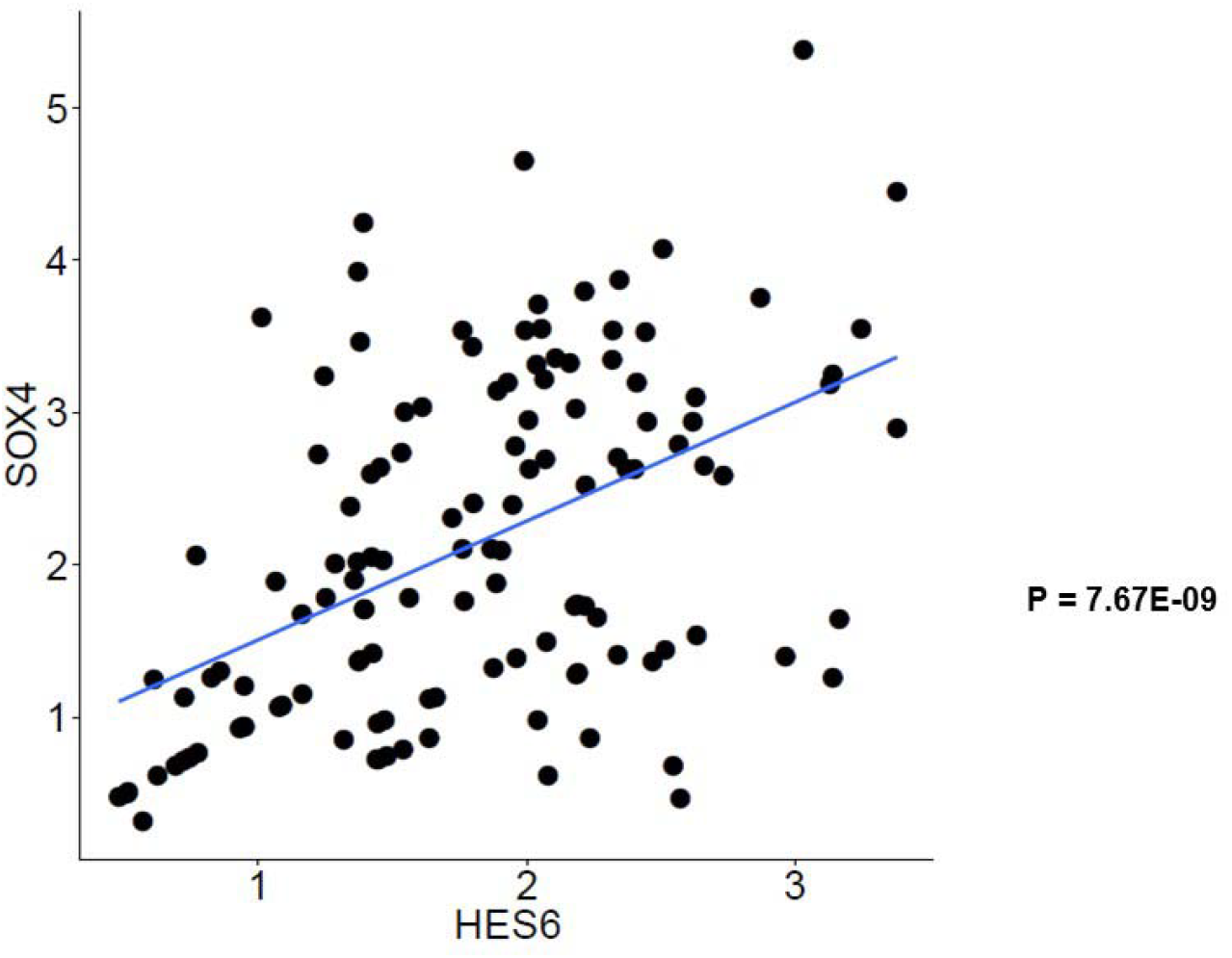
The co-expression of HES6 and SOX4 in individual goblet cells in the IM lesion.

**Supplementary Figure 14.**
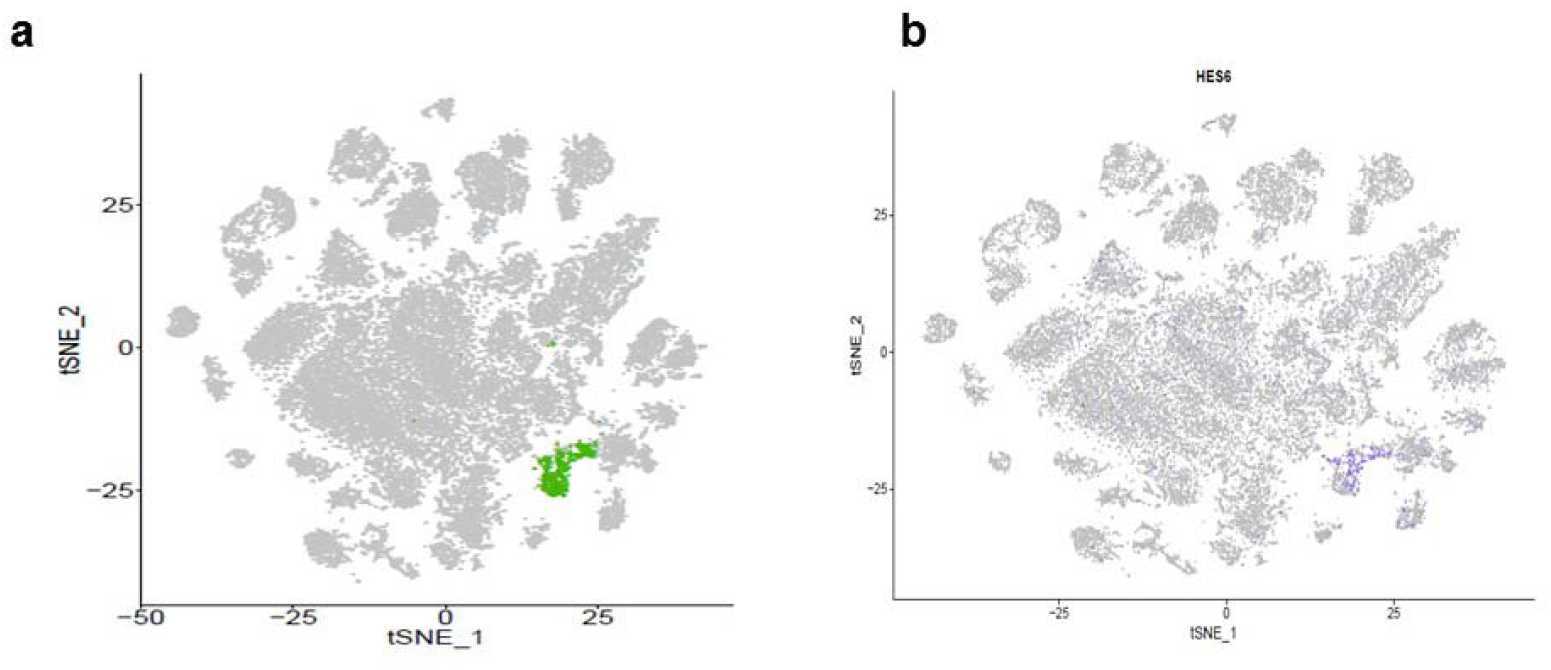
The t-SNE plot of the identified ‘goblet cell’ cluster (a) and the expression distribution of HES6 (b) (blue = high, gray = low).

**Supplementary Figure 15.**
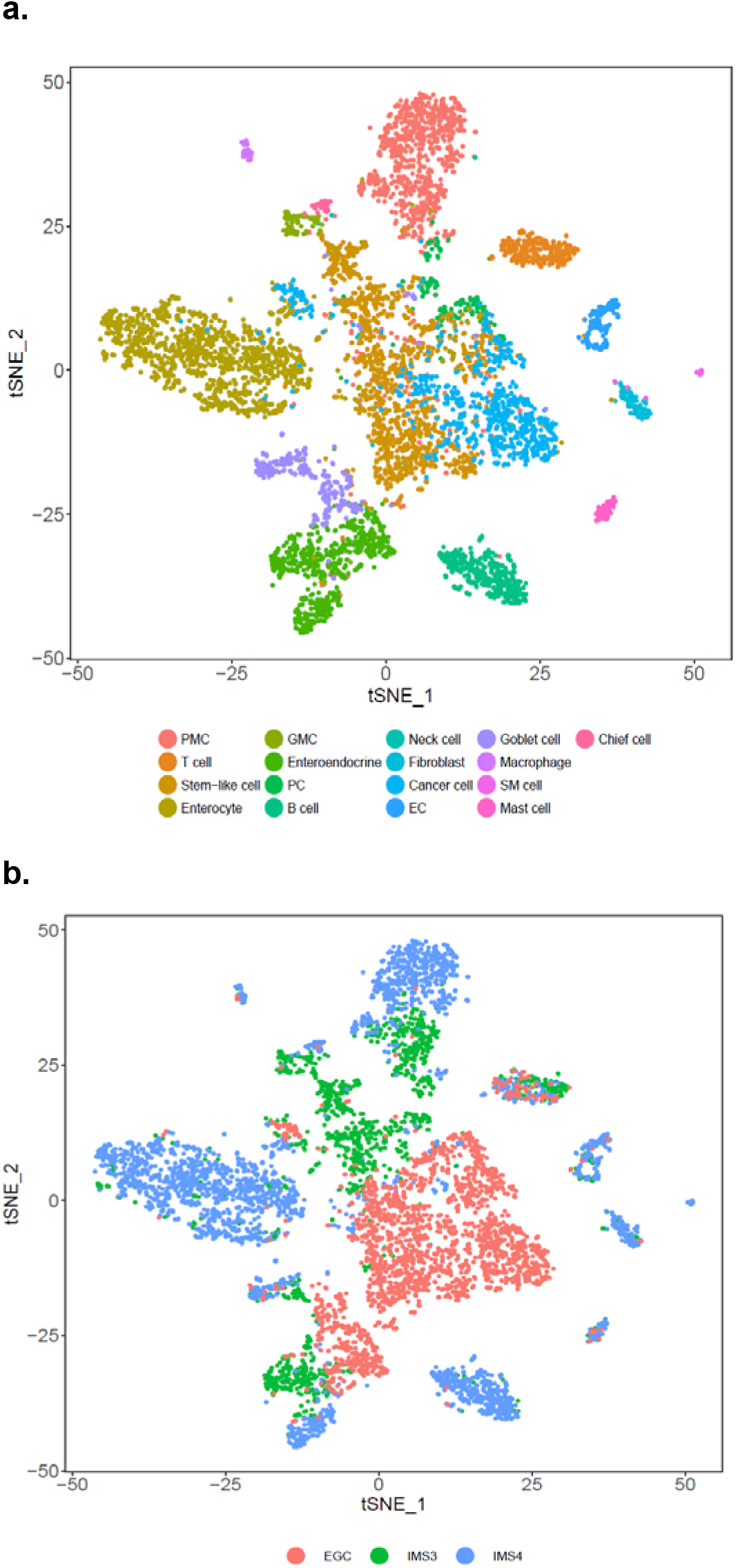
The t-SNE plot showing the distribution of diverse cell populations and biopsies in the same EGC patient (P8).

**Supplementary table 1.** The pathological grade of each biopsy used in the study.

| Sample ID | Histological diagnosis* | Patient |
| --- | --- | --- |
| NAG1 | NAG | P1 |
| NAG2 | NAG | P2 |
| CAG1 | CAG | P3 |
| CAG2 | CAG | P4 |
| CAG3 | CAG | P4 |
| IMW1 | IM-W | P5 |
| IMW2 | IM-W | P6 |
| IMS1 | IM-S | P7 |
| IMS2 | IM-S | P7 |
| IMS3 | IM-S | P8 |
| IMS4 | IM-S | P8 |
| EGC | EGC (High-grade dysplasia) | P8 |

\* NAG, Non Atrophic Gastritis CAG, Chronic Atrophic Gastritis, IM-W, Intestinal metaplasia with wild level; IM-S, Intestinal metaplasia with severe level; EGC, Early Gastric Cancer

**Supplementary table 2.** The clinical information of patients enrolled in the study.

| Patient ID | Age | Gender | Smoking<br>(cigs/day) | Alcohol<br>(units/week) | Dry<br>mouth | scorching<br>stomach<br>pain | family<br>history<br>of GC | H.P.<br>positive<br>(P vs N)* |
| --- | --- | --- | --- | --- | --- | --- | --- | --- |
| P1 | 58 | male | 0 | 0 | N | N | N | N |
| P2 | 56 | Female | 0 | 0 | Y | Y | N | P |
| P3 | 51 | male | 2 | 4 | N | N | N | P |
| P4 | 62 | Female | 0 | 0 | Y | N | N | N |
| P5 | 63 | male | 1 | 3 | N | N | N | P |
| P6 | 48 | Female | 0 | 0 | N | Y | N | N |
| P7 | 68 | male | 0 | 0 | Y | N | Y | P |
| P8 | 67 | male | 0 | 1 | Y | N | N | N |

\*P: *H.p* positive, N: *H.p* negative.

**Supplementary table 3.** The number of high-quality cells from each sample.

| Sample ID | NAG<br>1 | NAG<br>2 | CAG<br>1 | CAG<br>2 | CAG<br>3 | IMW<br>1 | IMW<br>2 | IMS<br>1 | IMS<br>2 | IMS<br>3 | IMS<br>4 | EGC |
| --- | --- | --- | --- | --- | --- | --- | --- | --- | --- | --- | --- | --- |
| # of cells | 2030 | 1627 | 2988 | 6570 | 4827 | 131<br>5 | 125<br>3 | 219<br>8 | 178<br>8 | 156<br>4 | 278<br>4 | 222<br>0 |

**Supplementary table 4.** The known markers for cell lineages in stomach.

| Categories | Cell lineages | Marker genes |
| --- | --- | --- |
| Endocrine | G cell | GAST |
|  | X cell | GHRL |
|  | D cell | SST |
| Mucous and secretory lineages | pit mucous cell (PMC) | MUC5AC |
|  | gland mucous cell (GMC) | MUC6 |
|  | parietal cell | ATP4A,ATP4B,GIF |
|  | chief cell | PGA4,PGA3,LIPF |

**Supplementary table 5.** The marker genes of each identified cell type in the atlas.

|  |  |  |
| --- | --- | --- |
| Immune cells | T cell | CD2, CD3D,CD3E,CD3G |
|  | B cell | CD79A,CD19 |
|  | mast cells | TPSAB1,TPSB2 |
|  | Macrophage | CD14, CD163, CD68, CSF1R |
| Stromal cells | Fibroblasts | FAP, PDPN,COL1A2,DCN,<br>COL3A1, COL6A1 |
|  | Endothelial cells | PECAM1,VWF,ENG,MCAM |
| Stem cell | stem cell | OLFM4,SOX2,LGR5,CCKBR |
| Myocytes | Smooth muscle cell (SMC) | ACTA2,ACTN2,MYL2,MYH2 |
| Proliferative cell | proliferative cell (PC) | MKI67,BIRC5,CDK1 |
| Intestinal cells | goblet cell | TFF3,REG4,MUC2 |
|  | enteroendocrine cell | CHGA,CHGB,TAC1,TPH1,NEU<br>ROG3 |
|  | enterocytes | FABP1,CA1,VIL1 |

**Supplementary table 6.** Gene signatures for each epithelial cell type in each lesion and enriched pathways for the gene signature of each lesion.

**Supplementary table 7.** Genes showing differential expression between pit mucous cells and gland mucous cells, and their enriched pathways.

**Supplementary table 8.** Genes that differentially expressed between goblet subsets and their enriched pathways.

**Supplementary table 9.** The pathways enriched for cancer cell-elevated genes.

**Supplementary table 10.** The dysregulation of cancer cell-elevated genes in TCGA.

***Supplementary table 5-10 are provided in the Extended Datasheet.***

